# Post-eclosion growth in the *Drosophila* Ejaculatory Duct is driven by Juvenile Hormone signaling and is essential for male fertility

**DOI:** 10.1101/2024.08.12.607650

**Authors:** Navyashree A. Ramesh, Allison M. Box, Laura A. Buttitta

## Abstract

The *Drosophila* Ejaculatory duct (ED) is a secretory tissue of the somatic male reproductive system. The ED is involved in the secretion of seminal fluid components and ED-specific antimicrobial peptides that aid in fertility and the female post-mating response. The ED is composed of secretory epithelial cells surrounded by a layer of innervated contractile muscle. The ED grows in young adult males during the first 24h post-eclosion, but the cell cycle status of the ED secretory cells and the role of post-eclosion ED growth have been unexplored. Here, we show that secretory cells of the adult *Drosophila* ED undergo variant cell cycles lacking mitosis called the endocycle, that lead to an increase in the cell and organ size of the ED post eclosion. The cells largely exit the endocycle by day 3 of adulthood, when the growth of the ED ceases, resulting in a tissue containing cells of ploidies ranging from 8C-32C. The size of the ED directly correlates with the ploidy of the secretory cells, with additional ectopic endocycles increasing organ size. When endoreplication is compromised in ED secretory cells, it leads to reduced organ size, reduced protein synthesis and compromised fertility. We provide evidence that the growth and endocycling in the young adult male ED is dependent on Juvenile hormone (JH) signaling and we suggest that hormone-induced early adult endocycling is required for optimal fertility and function of the ED tissue. We propose to use the ED as a post-mitotic tissue model to study the role of polyploidy in regulating secretory tissue growth and function.

## Introduction

The Drosophila male reproductive tract consists of a pair of sperm-producing testes connected to seminal vesicles, which along with a pair of accessory gland lobes, empty into an organ called the Ejaculatory Duct (ED). The ED is a site of sperm and seminal fluid mixing prior to transfer to the Ejaculatory bulb for female insemination during mating (Avila et al., 2016; Bertram et al., 1992; Hopkins et al., 2019) and these secretions are transferred into the ejaculatory duct during copulation through a transition zone called the anterior ED papillae (Bairati, 1968). ED cells are derived from the portion of the genital disc arising from the A9 primordium, and thus are ectodermal in origin (Ahmad and Baker, 2002; Avila et al., 2016) The ED also produces seminal fluid components, and by proteomic analysis of ED tissue, 138 peptides have been identified, of which 63 were proposed to be seminal fluid proteins (SFPs) (Sturm et al., 2021; Takemori and Yamamoto, 2009). Peptides secreted by the ED are involved in processes of lipid modification, immune response, catalysis, hormone-like activity, sperm storage, cell adhesion, odorant-binding proteins, and cellular metabolism (Takemori and Yamamoto, 2009). Contents from the ED are transferred to the Ejaculatory bulb and then to the female reproductive system via contractions of muscle, which surround the ED (Avila et al., 2016; Lung et al., 2001; Wilson et al., 2017). ED-produced antimicrobial peptides such as Cecropin A1 and the ED-specific Andropin (Anp) may increase fertility by reducing infections upon mating (Date-Ito et al., 2002; Lung et al., 2001; Samakovlis et al., 1991). Other SFPs are thought to promote male fertility through multiple mechanisms that impact sperm storage and female post-mating response. For example, the ED produces a pheromone peptide termed Ductus ejaculatorius peptide (Dup) Dup99B, which has homology to Sex Peptide (Accessory gland protein, Acp70), and like Sex Peptide, can elicit the female post-mating responses of increased egg laying and reduced sexual receptivity (Rexhepaj et al., 2003; Saudan et al., 2002). The anterior ED produces Esterase 6 (Est-6) a carboxylesterase transferred to females during copulation, which is a pheromone and odorant degrading enzyme that influences female sperm storage, sperm usage and re-mating (Chertemps et al., 2012; Ludwig et al., 1993; Richmond et al., 1980; Saad et al., 1994). Esterase-6 is also involved in the synthesis of sex pheromones (Takemori and Yamamoto, 2009). The ED also produces Glucose dehydrogenase (Gld) an enzyme that aids in sperm storage in the female reproductive tract after mating (Cavener, 1985; Cox-Foster et al., 1990; Iida and Cavener, 2004). The ED is therefore an important tissue of the male reproductive system that provides components to optimize male fertility, in addition to serving a structural role in transferring seminal fluid.

Despite important predicted functions for the ED in males, little is known about the development and growth of this tissue. Flies lacking the protein tyrosine kinase 7 Wnt co-receptors *off-track* (*otk*) and *off-track 2* (*otk2*) are viable but were reported to be male sterile due to defective morphogenesis of the ED (Linnemannstons et al., 2014). In these mutants, it is unclear whether the ED defects leading to infertility were structural due to defects in the muscle layer, or defects in the secretory cells, or both. Here, we examine the post-eclosion growth and cell cycle dynamics in the secretory epithelium of the ED. We show that this tissue undergoes additional cell cycles lacking M-phase that increase cellular ploidy, resulting in increased cell and tissue size, which supports a high level of protein synthesis that promotes male fertility. Post-eclosion endocycles in the ED are conserved in different species of *Drosophila,* and we show that compromising early adult endocycles in the ED reduces tissue growth, protein synthesis and male fertility. Juvenile hormone (JH) signaling is required for reproduction in Drosophila males (Baumann et al., 2017; Kurogi et al., 2024; Zhang et al., 2021), and here we show an essential role for JH signaling in promoting early post-eclosion endocycling of the secretory cells of the ED.

## Results

### Identification of cell types and their organization in the adult *Drosophila* Ejaculatory duct

The male adult *Drosophila* Ejaculatory duct is known to be involved in peristaltic motion to transfer sperm from the testes and seminal vesicles as well as seminal fluid containing fertility factors, enzymes, hormones and antimicrobial peptides to females (Avila et al., 2016; Bertram et al., 1992; Lung et al., 2001; Norville et al., 2010; Wilson et al., 2017). Little is known about the cell types in this tissue and their organization. To explore the different cell types in the ED we stained the tissue with cell type-specific antibodies to examine epithelial cells, neurons and muscle. Similar to a previous report (Susic-Jung et al., 2012), we observed that the ED epithelium is surrounded basally with striated muscles revealed by actin and myosin staining as well as staining with the muscle cell-specific marker Tinman (Liu et al., 2009; Zaffran et al., 2006)(Supp. Figure 1B-C). We observed a monolayer of secretory epithelial cells with larger nuclei expressing the epithelial cell-cell junction marker Fasciclin III (FasIII), forming a lumen apically (Supp. Figure 1C’’). The basal surface of these large epithelial cells contains Integrin β_ps_, an extracellular matrix receptor (Supp. Figure 1D). The peristaltic contraction of the ED suggests the muscle should be innervated. We identified the ED neurons using the neuronal markers Fustch and Elav (Supp. Figure 1E,F). This revealed the neuronal network overlying the basal muscle layer and labeled the smaller diploid nuclei of the ED. We noted that all diploid nuclei we could find using DAPI labeling in the ED are Elav positive, suggesting the muscle nuclei also express Elav (Supp. Fig. 1F). This tissue organization demonstrates the secretory ED epithelium contains enlarged cells forming a lumen surrounded by an ECM layer attached to innervated muscle.

### Endoreplication of secretory cells of the adult *Drosophila* Ejaculatory duct leads to post-eclosion organ growth

We recently described a post-eclosion wave of endocycling that occurs in the male accessory gland that promotes gland growth and function in male fertility (Box et al., 2024). During those studies, we also observed that the ED grows post-eclosion. We measured the area of ED on the day of eclosion (DOE) and at several time points post-eclosion (PE). The area of the ED increases until day 5 and then remains constant without any further increases all the way to day 55 of adulthood (Figure 1 A,B, Supp. Figure 2A). We also observed that the area of ED secretory cell nuclei significantly increased by day 10 PE compared to DOE (Figure 1C). To assay for DNA replication/synthesis we labeled live tissues with the S-phase DNA synthesis marker 5-Ethynyl-2′-deoxyuridine (EdU) (Flomerfelt and Gress, 2016). We observed widespread DNA replication on the day of eclosion in this tissue, which we also confirmed with EdU feeding assays in vivo (Figure 1 D). However, we did not observe any evidence of cells undergoing mitosis (Figure 1D). To ensure we did not miss any mitotic events, we stained adult EDs with the mitotic marker phospho-serine 10 Histone H3 (PH3) and labeled *ex-vivo* with EdU at one-hour intervals throughout the day of eclosion. We observed DNA synthesis during each hour across 24h, but no mitotic labeling (Supp. Fig 2C). These results suggest that the secretory cells of adult *Drosophila* ED are postmitotic but not quiescent and enter an endocycle.

**Figure 1:**
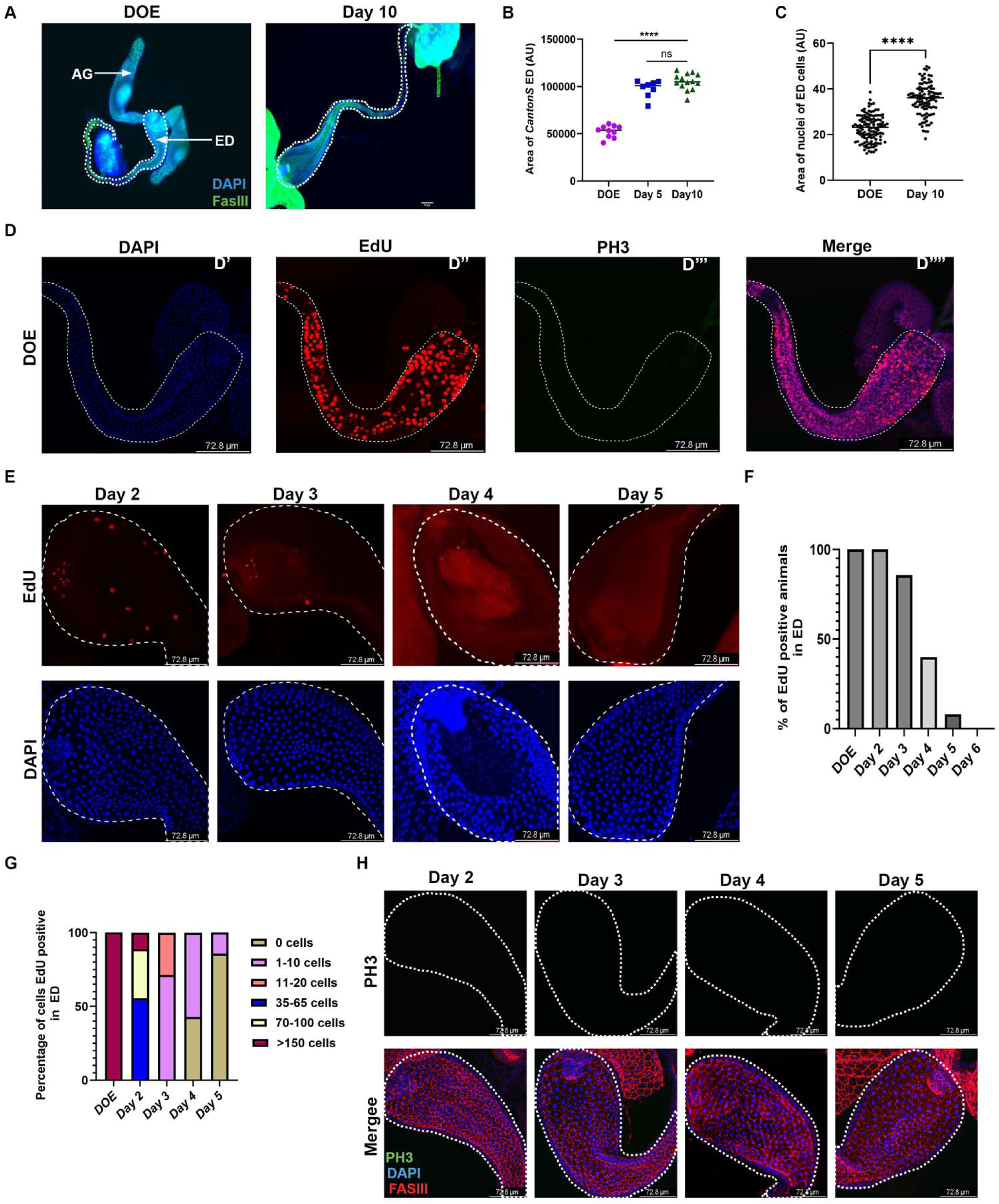
Endoreplication of secretory cells of the adult Drosophila Ejaculatory duct and post-eclosion organ growth. A. Images of adult ejaculatory duct (ED) on the day of eclosion (DOE) and day 10 showing an increase in the tissue size on Day 10. The ED is outlined with a dotted line. The tissue is labeled with Fas III antibody. The images were captured at 10X magnification. Scale bar: 70µm B. Quantification of the area of ED on DOE, Day 5, and Day 10 post-eclosion shows that the size of the ED increases from DOE to Day 5 and the growth ceases after Day 5 post-eclosion. Each dot represents the area of one ED from each male fly. C. Quantification of the area of the nuclei of the secretory cells of ED on DOE and Day 10 post-eclosion showing the increase in nuclei size. D. Confocal micrographs of the ED on DOE (0-2hrs post-eclosion) with EdU incorporation in secretory cells of ED with no mitotic positive PH3 cells. The EdU assay in this panel was done with *ex vivo* soaking of the ED in EdU for 1 hour. D’-DAPI, D’’-EdU staining, D’’’-mitosis marker PH3 antibody staining. D’’’’-Merge with FasIII staining of ED. E. Confocal micrographs of ED with EdU incorporation that was done by feeding the adult animals with EdU for 24hrs prior to time points on Day 2, Day 3, Day 4, and Day 5. Nuclei were labeled with DAPI. F. Quantification of the percentage of males that are EdU positive on DOE, Day 2, Day 3, Day 4, and Day 5 post-eclosion. G. Quantification of the percentage of cells that are EdU positive in each ED on DOE, Day 2, Day 3, Day 4, and Day 5 post-eclosion. We classified the total EdU-positive cells into multiple groups: 0 cells, 1-10 cells, 11-20 cells, 35-65 cells, 70-100 cells, and>150 cells that are EdU-positive. H. Micrographs of ED stained with the mitotic marker Phosphohistone H3 Ser10 (PH3) on Day 2, Day 3, Day 4, and Day 5. We did not observe any cells undergoing mitosis at these time points. Statistical analysis performed for Figure 2B, C-Two-tailed unpaired t-test. P<0.0001 **** Scale bar for panels D, E, H – 72.8 µm.

To determine when the endocycle ceases in the adult ED, we performed EdU feeding assays at 24-hour intervals up to day 6 post eclosion. Endocycling of ED secretory cells slowly decreases from day 2 with almost no cells continuing to endocycle by day 5 post-eclosion (Figure 1E-G). We also looked for any mitotic events up to day 5 post eclosion but observed no evidence of cell proliferation (Figure 1H). This suggests the post eclosion growth of the ED is driven in part by endoreplication of the secretory cells.

### Post eclosion endoreplication and growth of the adult Ejaculatory duct secretory cells in different species of *Drosophila*

We recently showed that the post eclosion wave of endocycling in the accessory gland is conserved in other *Drosophila* species. During the course of those studies, we also discovered that the post-eclosion endoreplication of the secretory cells of the adult ED is conserved in other *Drosophila* species. *D. simulans,* and *D. yakuba,* are sister species of *melanogaster* (Markow and O’Grady, 2006), while *D. pseudoodscura* and *D. willistoni* belong to the *sophophora* group (Takashima et al., 2023). We observed post eclosion endocycling in all of these species, including the outgroup *D. virilis*, which exhibits slower developmental timing and therefore was assessed 16 hours post eclosion (Figure 2 A-C, Supp. Fig. 3A). Of note, the two species with the least post eclosion growth of the ED, *D. pseudoodscura* and *D. willistoni,* also exhibited the smallest number of endocycling cells post-eclosion (Figure 3 C, D). Post-eclosion endoreplication of the ED secretory cells is therefore conserved in *Drosophila* and correlates with post eclosion organ growth.

**Figure 2:**
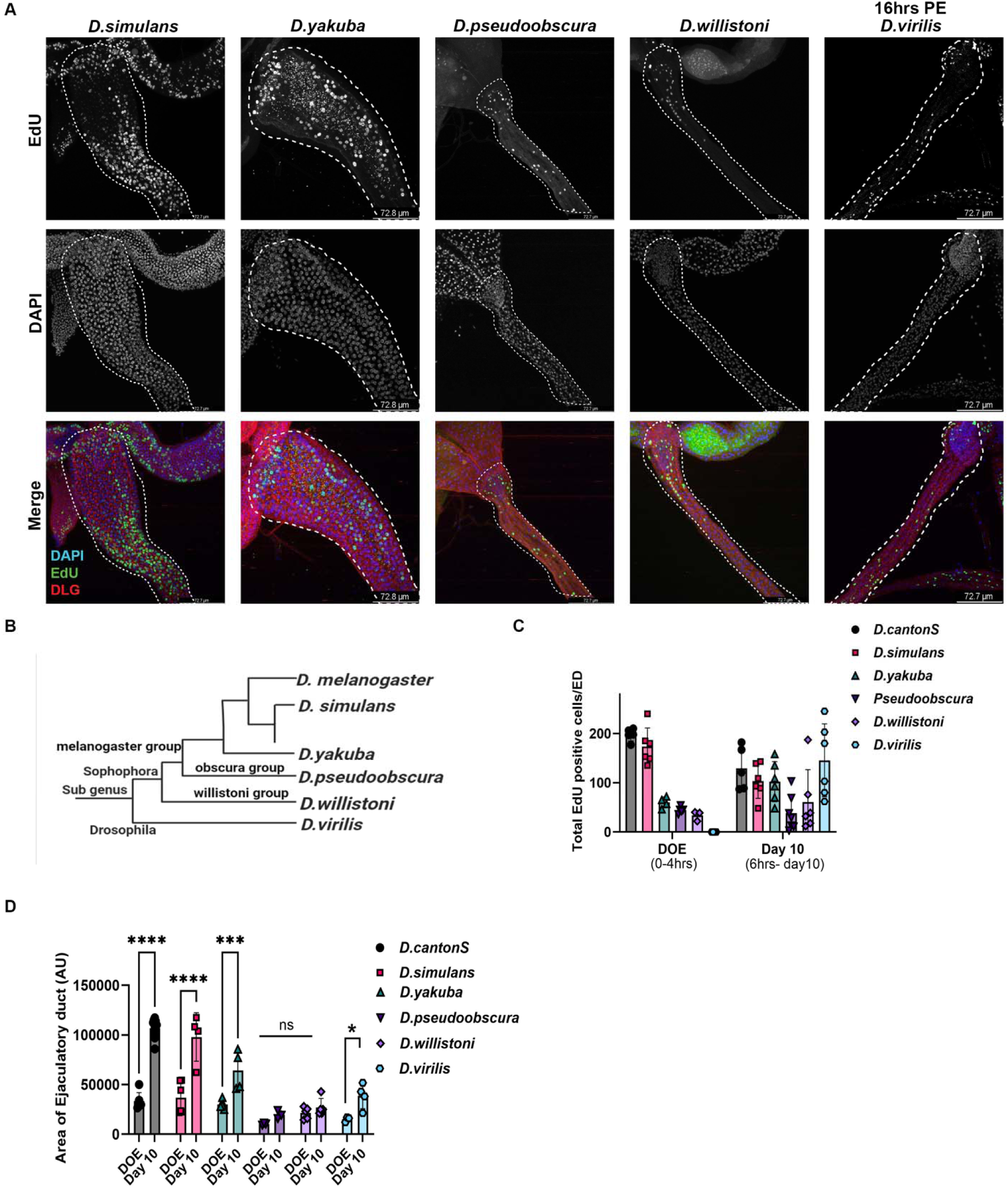
Post-eclosion endoreplication and growth of the ED is are conserved in different species of Drosophila. A. Confocal micrographs of EDs of different *Drosophila* species labeled with EdU on DOE (0-2hrs post-eclosion). This assay was performed by *ex vivo* incubation of the ED for 1 hour in EdU. The nuclei of secretory cells are labeled with DAPI, and the cell junctions of the secretory cells are labeled with anti-discs large (DLG). Scale bar:72.8 µm B. Phylogeny tree for subgroups of the different *Drosophila* species examined. C. Quantification of total EdU positive cells in different species on DOE performed with *ex vivo* EdU incorporation within 0-4 hrs post-eclosion. For day 10 EdU quantification we fed animals EdU until Day 10 and quantified the EdU incorporation on Day 10 (Male feeding begins ∼6h post eclosion). D. Quantification of the area of the ED on DOE (0-2hrs PE) and Day 10. Statistical test performed: Two-way ANOVA with multiple comparisons between DOE and Day 10 for respective species. P<0.0001 ****

**Figure 3:**
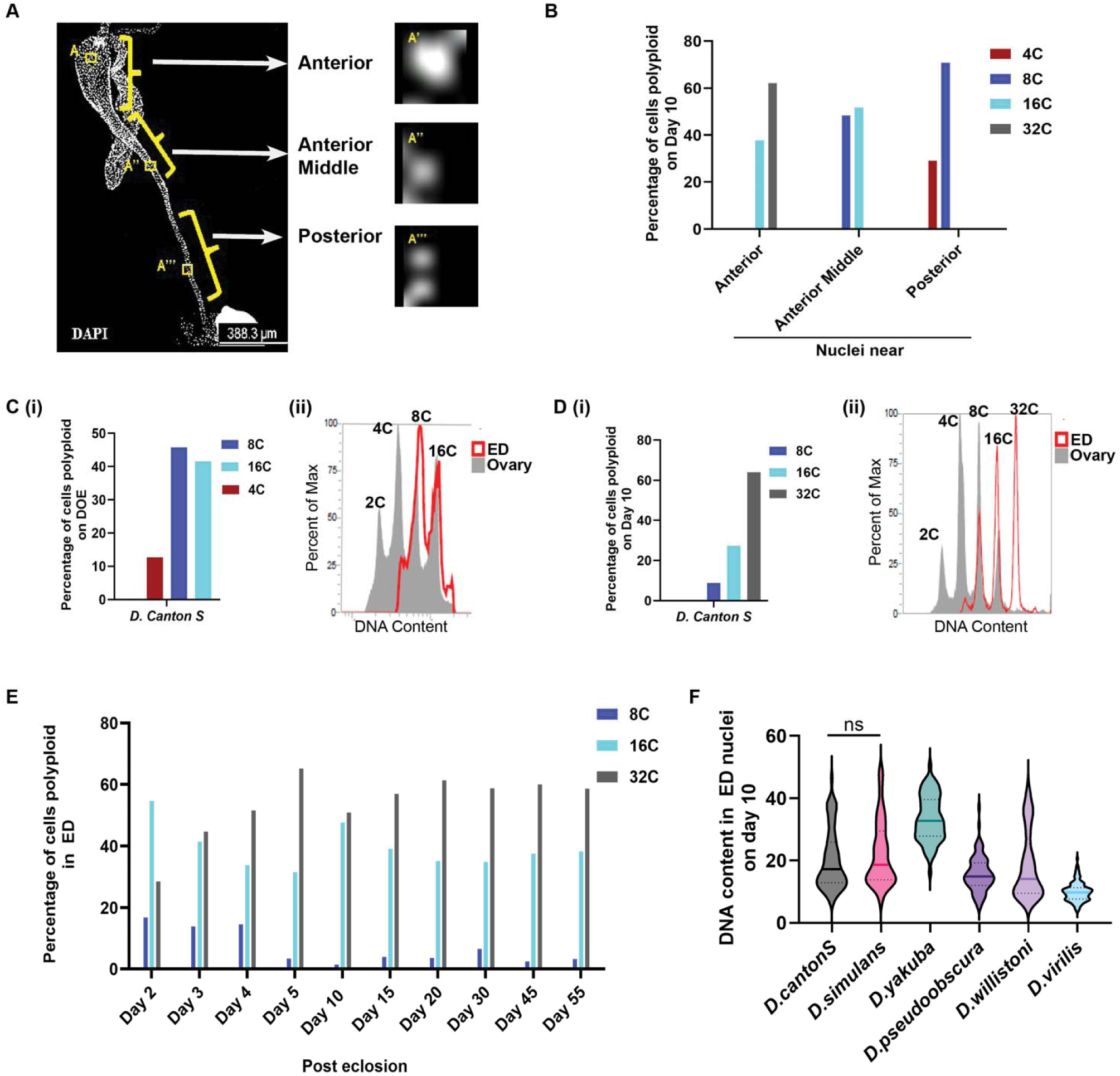
*Drosophila* Ejaculatory duct secretory cell ploidy correlates with organ shape and size. A. Image of a whole ED, showing that the volume of the duct reduces from anterior to posterior end. Based on this we classified ED secretory nuclei as Anterior, Anterior mid, and posterior. Zoomed images A’, A’’, A’’’ represents the nuclei of secretory cells from these respective regions of ED. Scale bar:388.3µm B. Quantification of the DNA content/ploidy from the nuclei of the secretory cells from the Anterior, Anterior middle, and posterior end regions of the ED. We quantified the ploidy from Day 10 old EDs by measuring the DAPI intensity and normalized it to the diploid cells from that specific region. C. (i) Quantification of the DNA content/ploidy on DOE by measuring the DAPI intensity, the endocycling secretory cells have ploidies of 4C, 8C, and 16C on DOE. (ii) Quantification of the ploidy of the secretory cells of ED by flow cytometry performed by isolating the nuclei from ED on DOE. The peak shaded in grey is Ovary and the peak with red color is from the ED. D. (i) Quantification of the DNA content/ploidy from the secretory cells of the ED on Day 10 post-eclosion by measuring the DAPI intensity. (ii) Quantification of the ploidy from the secretory cells of the ED by flow cytometry performed by isolating the nuclei from ED on Day 10 post-eclosion. The peak shaded in grey is Ovary and the peak with red color is from the ED. E. Quantification of the DNA content/ploidies of secretory cells of the ED performed by measuring the DAPI intensity on different days post-eclosion showing that the ploidy distribution is unchanged and constant after Day 5 PE. F. Quantification of the DNA content/ploidy of the secretory cells by measuring the DAPI intensity in different species, showing that the secretory cells of EDs of different *Drosophila* species we used are polyploid.

### *Drosophila* ED secretory cell ploidy correlates with organ shape and size

Recent work in the *Drosophila* heart demonstrated that cellular ploidy and regulation of endoreplication can influence tissue morphogenesis and function in a contractile organ (Chakraborty et al., 2023). The anterior portion of the ED where the accessory glands and seminal vesicles meet with the anterior ED papillae is larger and the organ tapers and thins towards the posterior end, where it connects to the ejaculatory bulb (Figure 3A). We therefore measured nuclear areas in the anterior, middle and posterior regions of the mature tissue on day 10, and observed that nuclei towards the anterior are larger compared to the nuclei of the secretory cells towards the anterior mid and posterior end as shown in Figure 3A. We further confirmed the differences in ploidy by quantifying integrated DAPI intensities for the nuclei of the secretory cells, normalized to the diploid Elav+ cells of the organ (Figure 3B). The cells in the anterior region have a DNA content of 16C and 32C, while the cells in the anterior middle have a DNA content of 8C and 16C, and the cells in the posterior end have a DNA content of 4C and 8C (Figure 3B). These different regions of ED may be involved in providing different secretions or different amounts of secretions (Sturm et al., 2021). Thus, like the *Drosophila* heart, the ED exhibits regional-specific polyploidy of the tissue, which correlates with the tissue size and organ morphogenesis.

To confirm our image-based ploidy measurements and understand how these ploidies arise, we developed a flow cytometry assay to isolate nuclei from the adult EDs and measure DNA content by flow cytometry at timepoints post-eclosion. As a control we used ovaries of age and genotype matched females, which have a known ploidy distribution and serve as a control to accurately assign DNA content peaks (Bosco et al., 2007). On the day of eclosion, ED secretory cells exhibit ploidies of primarily 8C and 16C (Figure 3C), with some intermediate ploidies consistent with their replicating EdU+ status. By day 10, we observed a large population with 32C DNA content, as well as populations with 16C, and 8C (Figure 3D). Consistent with their EdU-negative non-cycling status, by day 10, we observe very few nuclei with intermediate ploidies. With the flow cytometry confirmation of our DAPI imaging measurements, we quantified the ED ploidy distribution using DAPI imaging in adults from young males (day 2 post eclosion) to aged males (day 55) (Figure 3E). We observed consistent ploidy measurements from day 15-55 in aged *D. melanogaster* virgin males suggesting additional endoreplication in aged virgin adults is rare under normal physiological conditions.

We noted differences in ED endocycling across *Drosophila* species correlated with post eclosion ED growth. However, our longer-term EdU labeling for 10 days revealed differences in nuclear areas across species (Supp. Figure 3B-D). To examine this in more detail we performed flow cytometry measurements of *D. simulans*, *D.* y*akuba* and *D. virilis* at day 10 using the species-matched female ovary to normalize DNA content measurements to the species genome size (Bosco et al., 2007) (Supp. Figure 3 E-G). Consistent with our EdU labeling and DAPI imaging quantifications (Figure 3F) we observe higher ploidies in *D. simulans* and *D. yakuba* compared with *D. virilis*. We also observe more cells of lower ploidies (<16C or less) in species with smaller, thinner EDs such as *D. pseudoobscura*, *D. willistoni* and *D. virilis* (Figure 3F, Supp. Figure 3B).

### Compromising the endoreplication of ED secretory cells results in reduced ED size and fertility

To manipulate the secretory cells of the ED, we performed a Gal4 screen to identify an ED-Gal4 driver. We screened through ∼100 Gal4 lines predicted to express in the male accessory glands and/or ED based on RNAseq data and identified Luna-Gal4 (InSite line BDSC 63746) as specific to the ED in the male reproductive system. We verified the Gal4 expression in the polyploid secretory cells of the ED using a Gal-4 responsive *U*pstream *A*ctivating *Sequence* (*UAS*) driving a nuclear localized RFP (*UAS-RFPnls*). We did not observe any expression in the muscle or neuronal layer of the ED, or any other parts of the male reproductive system including the accessory glands and ejaculatory bulb (Figure 4A, Supp. Figure 2F). We did not find the expression of Luna-Gal4 in any other adult *Drosophila* tissues except cardia (Supp. Figure 2F). Luna-Gal4 expression in the ED begins prior to eclosion (96h APF) and continues into adulthood. By combining the *Luna-Gal4,* with a *UAS-GFPnls* with a temperature-sensitive Gal80 that represses Gal4 (*Gal80^ts^*) (McGuire et al., 2004) we used temperature shifts to limit *Luna-Gal4* expression to the adult stages and confirmed that *Luna Gal4* continues to express Gal4 in the secretory cells of ED throughout adulthood, with and without mating, until day 70 post-eclosion (Supp. Figure 4A).

**Figure 4:**
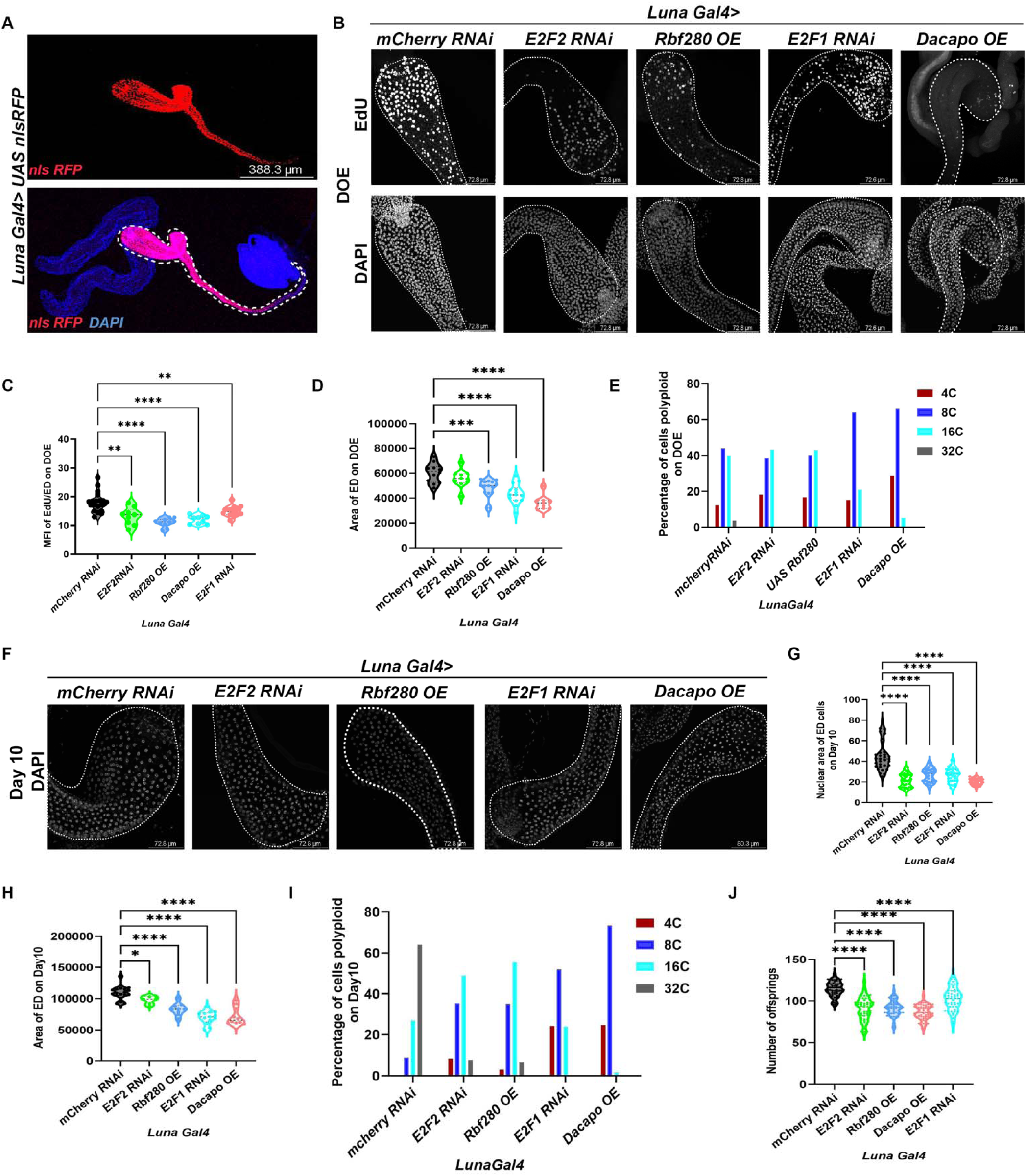
Compromising the endoreplication of ED secretory cells results in reduced ED size and fertility. A. Confocal micrograph of *Drosophila* male reproductive system showing the expression of *Luna Gal4* specifically in ED secretory cells verified with the expression of *UAS-nls-RFP* in the ED cells. Scale bar: 388.3µm B. Confocal micrographs of ED on DOE with knockdown of *E2F1, E2F2* using RNAi, Overexpression of *Rbf280,* and *Dacapo-GFP* using *Luna Gal4*. The upper panel is the EdU labeling on DOE with *ex-vivo* incubation in the background of all these manipulations in ED secretory cells. We stained the nuclei with DAPI to measure the DNA content. Scale bar:72.8 µm C. Quantification of the Mean fluorescence intensity of EdU on DOE with knockdown of *E2F1, E2F2* using RNAi, Overexpression of *Rbf280*, and *dacapo GFP* using *Luna Gal4*. D. Quantification of the area of the ED on DOE with knockdown of *E2F1*, and *E2F2* using RNAi, Overexpression of *Rbf280*, and *Dacapo-GFP* using *Luna Gal4*. E. Quantification of the DNA content/ploidy on DOE measured using the DAPI intensities in all the different backgrounds of Overexpression and knockdowns in secretory cells. F. Confocal micrographs of the ED on Day 10 post-eclosion labeled with DAPI show that the nuclei are smaller in *E2F1 RNAi, E2F2 RNAi, Rbf 280*, and *Dacapo-GFP* overexpression. Scale bar:72.8 µm G. Quantification of the nuclear area of the secretory cells of the ED on Day 10 post-eclosion showing reduced nuclear size in *E2F1 RNAi, E2F2 RNAi, Rbf 280*, and *Dacapo-GFP* overexpression conditions. H. Quantification of the Area of the ED on Day 10 post-eclosion showing reduced area of ED in *E2F1* RNAi, *E2F2* RNAi, *Rbf 280*, and *Dacapo-GFP* overexpression conditions. I. Quantification of the DNA content/ploidy of the secretory cells of the ED on Day 10 post-eclosion showing reduced ploidy in *E2F1* RNAi, *E2F2* RNAi, *Rbf 280*, and *Dacapo-GFP* overexpression conditions compared to the control. J. Quantification of fertility for males with reduced DNA content in the ED. Statistical Analysis: Two-way ANOVA with multiple comparisons between control and Different knockdown and overexpression conditions. P<0.0001 ****

We next used the *Luna-Gal4* driver to express transgenes to inhibit or slow the endocycle in the ED. Oscillations of the core cell cycle transcription factor complex *E2F* have been shown to drive the endocycle (Weng et al., 2003; Zielke et al., 2011), therefore we knocked down the two *Drosophila E2F* transcription factors *E2F1* and *E2F2* or overexpressed a constitutively active form of the E2F repressor, the Retinoblastoma family protein (*Rbf^280^*) (Xin et al., 2002) to limit S-phase entry and performed EdU labeling on the day of eclosion. All three approaches effectively reduced EdU labeling and nuclear ploidy in the ED on the day of eclosion, demonstrating they reduced or slowed post-eclosion S-phases. *Rbf^280^* expression and *E2F1* RNAi also effectively reduced adult ED size on the day of eclosion (Figure 4 B-E). We next overexpressed the G1-S *CyclinE/Cdk2* inhibitor *Dacapo*, which acts to limit G1/S entry and therefore also slows or blocks endoreplication when expressed at high levels (Swanson et al., 2015). As expected, *Dacapo* overexpression severely limited endocycling in the ED on the day of eclosion, resulting in reduced nuclear ploidies and reduced organ size (Figure 4 B-E).

We next examined tissues on day 10, several days after ED cells normally have ceased endocycling. Consistent with the reduced or blocked endocycling observed on the day of eclosion, by day 10 the EDs with inhibited endocycling exhibit reduced organ size, reduced nuclear size and reduced ploidies, demonstrating that these genetic cell cycle manipulations do not simply delay or slow the endocycle but instead effectively limit it (Figure 4F-I, Supp. Figure 4B). These results strongly suggest that the early endoreplication in secretory cells of ED post-eclosion is required for the growth of the ED. To test whether reduced ED size and ploidy may impact fertility, we crossed the males with reduced or inhibited endocycling to wild-type *Canton-S* females at a male-to-female ratio of 1:8 for individual male fertility assays. We observed significantly reduced offspring from males with reduced ploidy in the ED (Figure 4J), suggesting the smaller organ size and reduced DNA content may lead to reduced secretion of essential ED-specific factors.

### Inducing extra endoreplication in the ED leads to organ hypertrophy

We next asked whether genetic manipulations expected to promote S-phase entry and increase or prolong endocycling would impact ED size and function. Overexpression of the G1-S *Cyclin/Cdk* complexes including *CyclinE/Cdk2* and *CyclinD/Cdk4* in *Drosophila* male accessory glands leads to increased endoreplication, as does knockdown of the *E2F* inhibitor *Rbf* with RNAi (Box et al., 2024). On the day of eclosion, we observed mild increases in EdU incorporation and organ size with these manipulations (Figure 5A-C), but by day 10 of adulthood we observed significant increases in organ size, nuclear area and ploidy (Figure 5 D-G, Supp. Figure 4 C,D), suggesting that growth and endocycling are already maximally activated on the day of eclosion and that our manipulations led to additional endocycles resulting in higher ploidies, with many cells containing 64C DNA content, a state normally never observed in the ED. These results were further confirmed by knocking down the DNA replication licensing inhibitor *Geminin* (Diffley, 2004), which also led to a small increase in nuclei with 64C DNA content (Figure 5 A-G). Despite the increased organ size and DNA content, we observed no significant increases in male fertility (Figure 5H).

**Figure 5:**
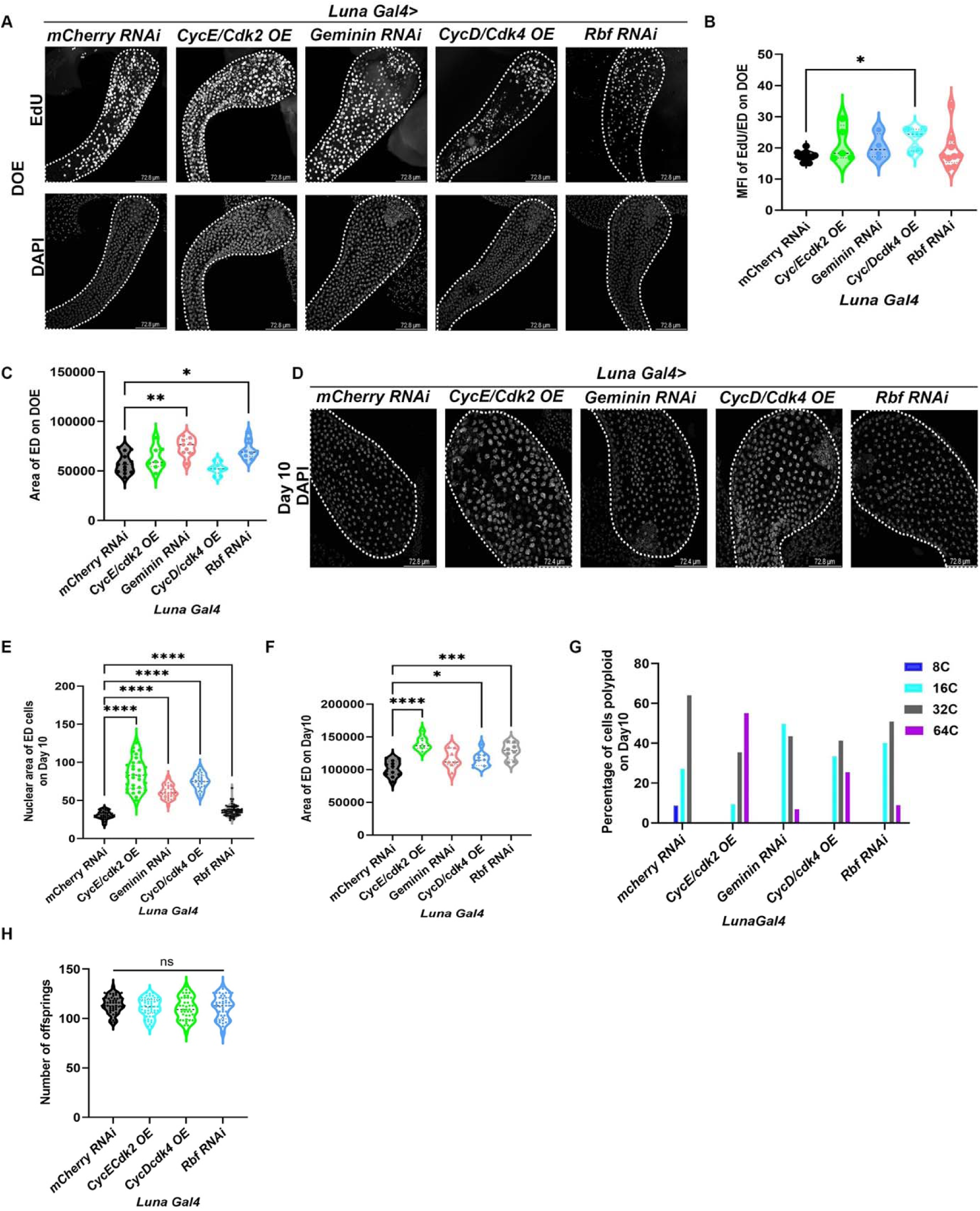
Inducing extra endoreplication in secretory cells of ED leads to organ hypertrophy. A. Confocal micrographs of EDs on DOE with overexpression of *CycE-Cdk2, CycD-Cdk4*, and knockdown of *Geminin, or Rbf* with RNAi using *Luna Gal4*. The upper panel is the EdU labeling performed by *ex-vivo* incubation on DOE, the lower panel is nuclei labeled with DAPI to measure the DNA content. Scale bar:72.8 µm B. Quantification of the Mean fluorescence intensity of EdU on DOE in the condition with overexpression of *CycE-Cdk2, CycD-Cdk4*, and *Geminin or Rbf* with RNAi driven by *Luna Gal4*. C. Quantification of the area of the ED with overexpression of *CycE-Cdk2, CycD-Cdk4,* and *Geminin or Rbf* with RNAi driven by *Luna Gal4*. D. Confocal micrographs of the ED on Day 10 post-eclosion labeled with DAPI show that the nuclei and the ED are larger with overexpression of *CycE-Cdk2, CycD-Cdk4,* and knockdown of *Geminin* or *Rbf* with RNAi using *Luna Gal4*. Scale bar:72.8 µm E. Quantification of the nuclear area of the secretory cells of ED on Day 10 post-eclosion with the indicated transgenes. F. Quantification of the area of the ED on Day 10 post-eclosion expressing the indicated transgenes. G. Quantification of the DNA content/ploidy of the secretory cells of ED on Day 10 post-eclosion using DAPI measurements showing increased ploidy with overexpression of *CycE-Cdk2, CycD-Cdk4*, and knockdown of *Geminin or Rbf* with RNAi using *Luna Gal4* compared to the control. H. Quantification of male fertility for animals expressing the indicated transgenes driven by *Luna-Gal4*. The animals with increased ED DNA content do not have significant differences in the number of offspring. Statistical Analysis: Two-way ANOVA with multiple comparisons between control and Different knockdown and overexpression conditions. P<0.0001 ****

### Stabilized Geminin effectively blocks adult endoreplication in the ED

Geminin is a DNA replication licensing inhibitor that is targeted for degradation during the cell cycle by the Anaphase Promoting Complex/Cyclosome (APC/C) ubiquitin ligase (Diffley, 2004; Zielke et al., 2008). A stabilized form of geminin lacking the “Destruction Box” site for APC/C-dependent ubiquitination dominantly blocks S-phase entry when overexpressed in *Drosophila* tissues. To rigorously test whether endocycling is required in the ED we expressed *stabilized Geminin* using the *Luna-Gal4* driver. EdU incorporation on the day of eclosion is nearly completely blocked by stabilized Geminin. EDs that normally exhibit >150 EdU+ nuclei exhibit at most 17 EdU+ nuclei when stabilized Geminin is expressed (Figure 6A-C). Blocking the endocycle led to a significant reduction in the area of the ED on DOE (Figure 6D) and a mild reduction in ploidy (Figure 6E). By day 10, EDs expressing *stabilized Geminin* still had not endocycled, indicating a nearly complete block, resulting in dramatically smaller organs (Figure 6F-G), cells with severely compromised ploidies (Figure 6 H, Supp. Figure 4E) and severely compromised male fertility (Figure 6I). This suggests the post eclosion endocycles in the ED are required for proper male organ function and fertility.

**Figure 6:**
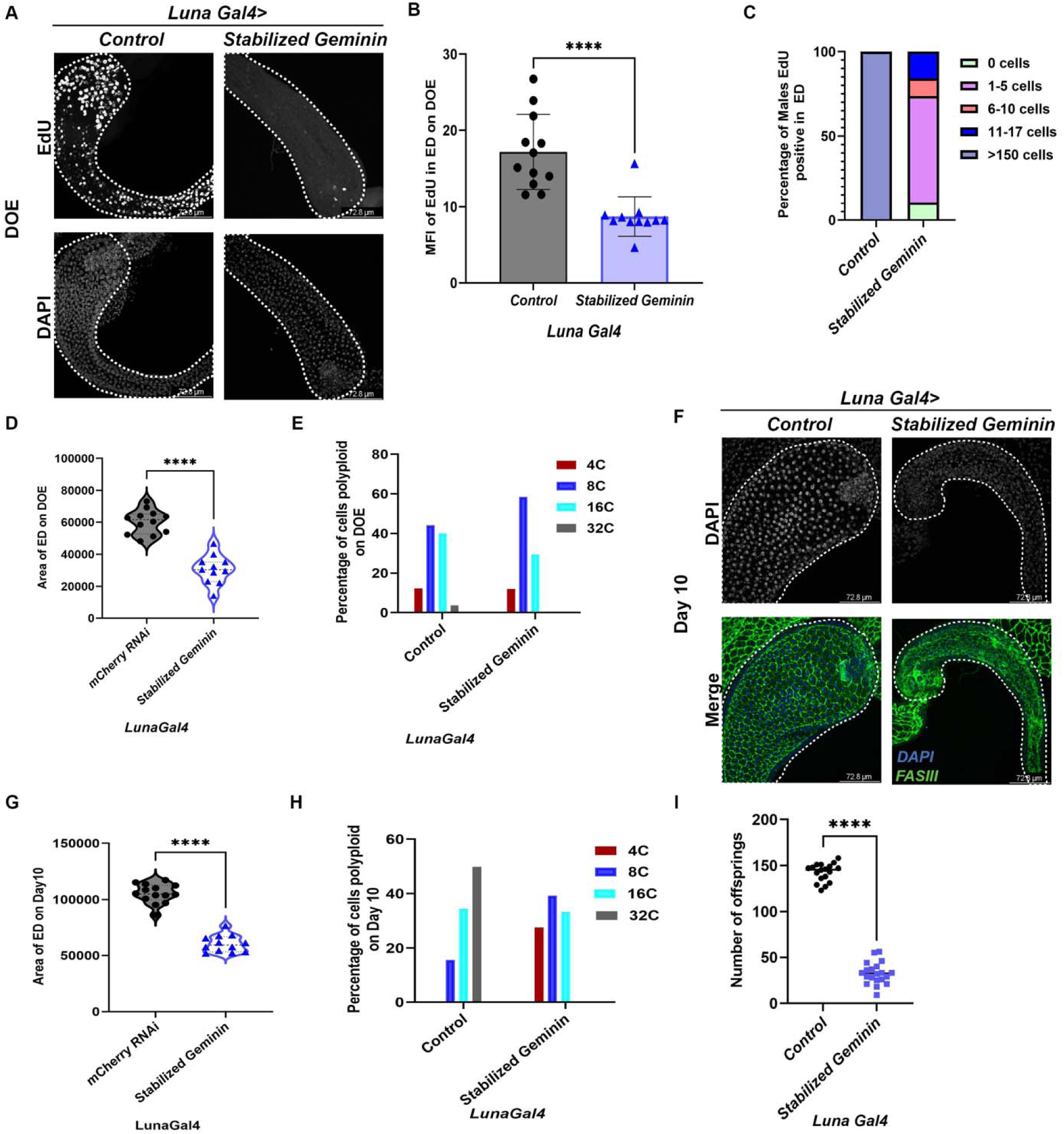
Stabilized Geminin effectively blocks the adult endoreplication in the ejaculatory duct. A. Confocal micrograph of the ED on DOE (0-2hrs) with EdU labeling performed ex vivo. A nearly complete block of endocycling is observed when stabilized *Geminin* is expressed. Nuclei are labeled with DAPI. Scale bar:72.8 µm B. Quantification of the mean fluorescence intensity of the EdU incorporated on DOE shows significantly reduced EdU with stabilized *Geminin* expression compared to the control. C. Quantification of the percentage of cells that are EdU positive on DOE in each ED performed with *ex vivo* EdU incorporation. We classified the total EdU-positive cells into multiple groups: 0 cells, 1-5 cells, 6-10 cells, 11-17 cells, and>150 cells that are EdU-positive in each ED of control and stabilized Geminin. D. Quantification of the area of ED on DOE with the indicated transgenes driven by *Luna Gal4*. E. Quantification of the DNA content/ploidy of the secretory cells on DOE with the indicated transgenes driven by *Luna Gal4*. F. Confocal micrographs of the ED on Day 10 post-eclosion labeled with DAPI and cell junction marker Fas III show that the nuclei are smaller with overexpression of stabilized *Geminin* compared to the control. G. Quantification of the area of ED on Day 10 with the indicated transgenes driven by *Luna Gal4*. H. Quantification of the DNA content/ploidy of the secretory cells of ED on Day 10 post-eclosion showing reduced ploidy in stabilized *Geminin* overexpression conditions compared to the control. I. Quantification of male fertility for animals with the indicated transgenes driven by *Luna Gal4*. The animals with *stabilized Geminin* overexpression have significantly reduced offspring compared to the control. Statistical Analysis: unpaired Two-tailed t-test. P<0.0001 ****

### Juvenile hormone (JH) signaling promotes post eclosion endoreplication and growth of the ED

JH is a family of acyclic sesquiterpenoids released by the endocrine gland Corpus allatum, which plays an important role in *Drosophila* development and reproduction (Flatt et al., 2005; Jindra et al., 2013). A pulse of JH is released immediately post-eclosion (Bownes and Rembold, 1987; Zhang et al., 2021) and in females JH has been shown to play an important role in inducing polyploidy in the fat body during vitellogenesis to promote reproduction (Ren et al., 2020; Riddiford, 2012; Wu et al., 2018; Wu et al., 2020). The well-studied nuclear effectors of JH signaling include *methoprene-tolerant* (*Met*), *germ-cell expressed* (*Gce*), and their co-factor *Taiman* (*Tai*) (Baumann et al., 2010; Li et al., 2019). We recently showed that JH receptors *Met* and *Gce* are required for post-eclosion endocycling in the accessory gland (Box et al., 2024). We therefore wondered whether this pathway may also mediate post-eclosion endocycling in the ED. We knocked down *Met, Gce*, and *Tai* via UAS-RNAi using *Luna-Gal4*. We observed reduced EdU incorporation on the day of eclosion in all three knockdowns compared to the control *mCherry* RNAi (Figure 7A, B). The area of the ED was also reduced on the day of eclosion and significant reductions in organ size and ploidy were observed at day 10 with JH receptor knockdowns, indicating the endocycles were not simply delayed (Figure 7 C-F). Male fertility was also reduced compared to the control when JH receptors were knocked-down (Figure 7G). These results suggest that JH signaling through *Gce*, *Met,* and *Tai* is required for post-eclosion ED growth via endocycles and male fertility.

**Figure 7:**
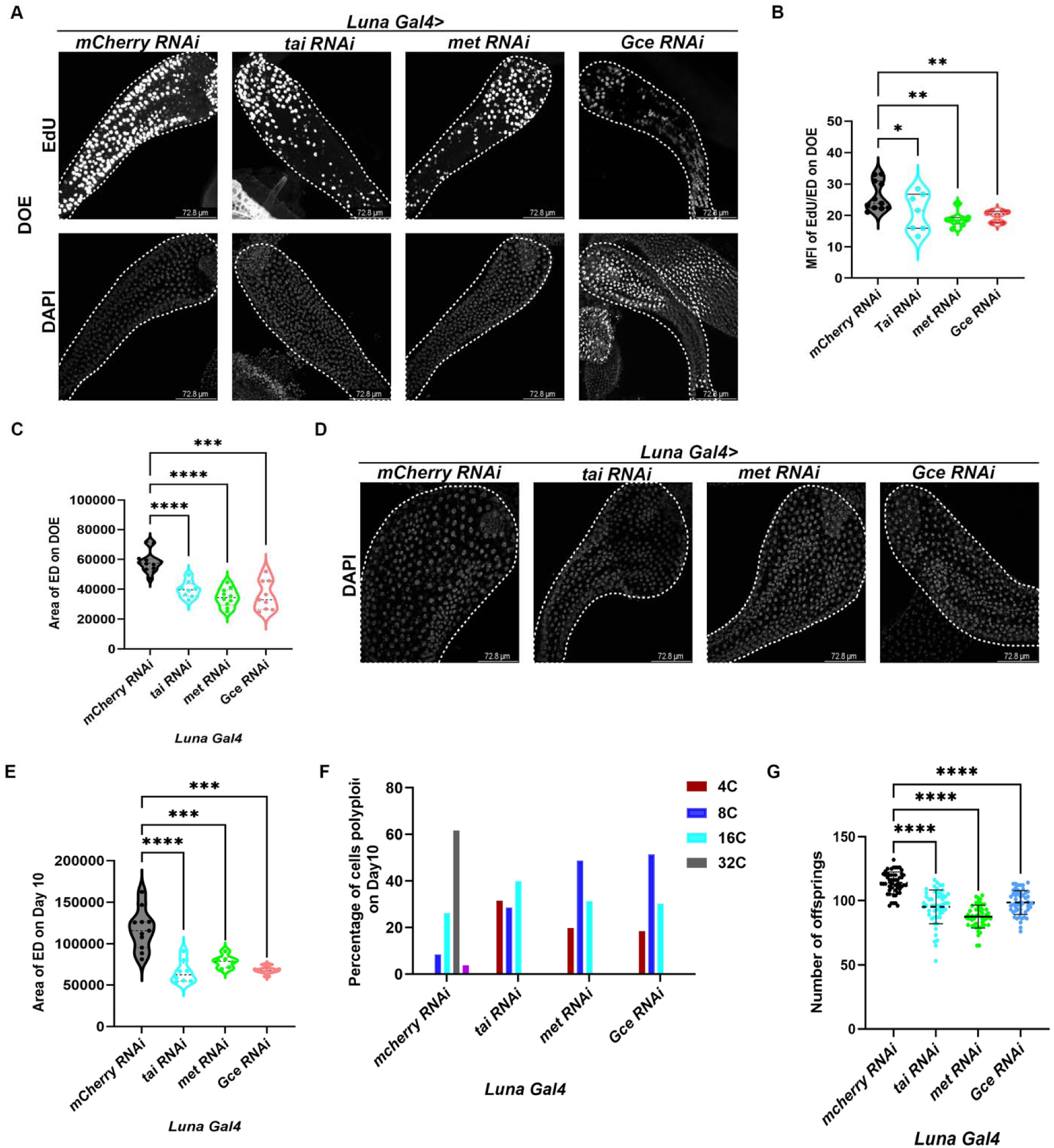
Juvenile hormone (JH) signaling promotes post-eclosion endoreplication and growth of *Drosophila* ejaculatory duct. A. Confocal micrographs of ED on DOE with knockdown of Juvenile hormone receptors *Gce, Met*, and transcription factor *tai* with RNAi using *Luna Gal4.* The control expresses *mCherry* RNAi driven by *Luna Gal4*. The upper panel is the EdU labeling on DOE. Nuclei are labeled with DAPI (lower panels) to measure the DNA content. Scale bar:72.8 µm B. Quantification of EdU incorporated on DOE shows reduced EdU with knockdown of *Gce*, *Met,* and *tai* with RNAi using *Luna Gal4* C. Confocal micrographs of the ED on Day 10 post-eclosion labeled with DAPI for the indicated transgenes driven by Luna Gal4. Nuclei are smaller with knockdown of *Gce, met, or tai* compared to the control. D. Quantification of the area of the ED on Day 10 post-eclosion for the indicated transgenes driven by Luna Gal4. E. Quantification of the DNA content/ploidy of the secretory cells of ED on Day 10 post-eclosion for the indicated transgenes driven by Luna Gal4. Reduced ploidy is observed for knockdown of *Gce, met or tai*. F. Quantification of male fertility for animals expressing the indicated transgenes driven by Luna Gal4. Fertility is reduced with the knockdown of *Gce, met or tai*. Statistical Analysis: Two-way ANOVA with multiple comparisons between control and Different knockdown and overexpression conditions. P<0.0001 ****

**Figure 8:**
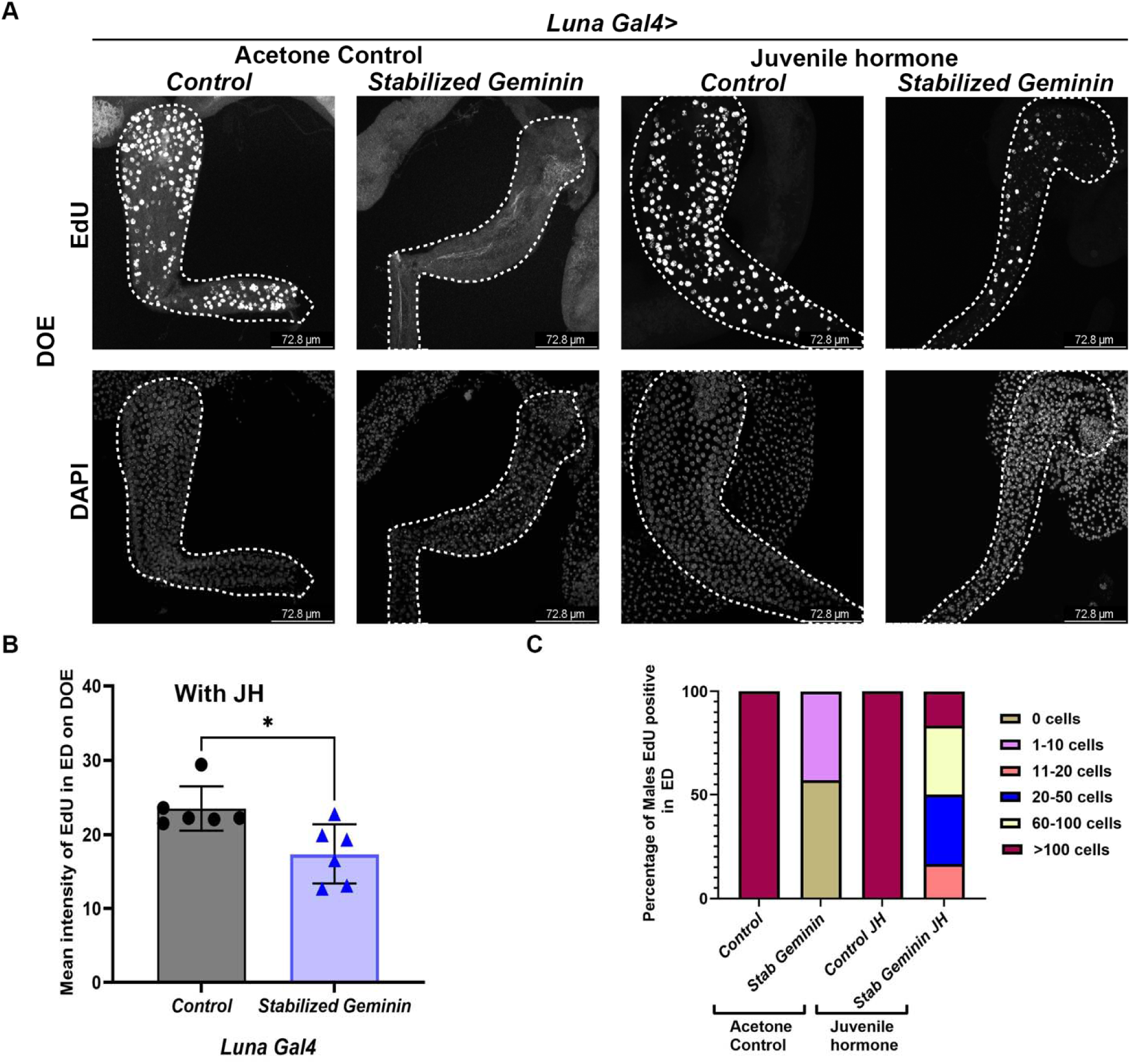
Juvenile hormone signaling is sufficient to induce endoreplication in the adult *Drosophila* ejaculatory duct. A. Confocal micrograph of EdU incorporation in ED on DOE (0-2 hrs PE) with vehicle control (Acetone)(A’, A’’) and Juvenile hormone (JH) III with 1-hour *ex-vivo* incubation(A’’’, A’’’’) along with the EdU in controls and EDs expressing stabilized *Geminin* to block the endocycle. Top panel EdU, bottom panel DAPI staining. Scale bar:72.8 µm B. Quantification of the Mean fluorescence intensity of the EdU per ED in both stabilized *Geminin* and controls incubated with JH III. C. Quantification of the percentage of cells that are EdU positive with *ex vivo* acetone and JH III incubation on DOE for EDs expressing the indicated transgenes. Controls express *Luna-Gal4* only. We classified the total EdU-positive cells into multiple groups: 0 cells, 1-10 cells, 11-20 cells, 20-50 cells, 60-100, and>150 cells that are EdU-positive in each ED of control and stabilized Geminin.

### Juvenile hormone signaling is sufficient to overcome blocked endoreplication in the adult ED

We recently showed that ectopic JH was sufficient to promote additional endocycling in post eclosion accessory glands (Box et al., 2024). We tested this in the ED, but the level of endocycling on the day of eclosion in the ED is already so high under control conditions, we found it difficult to assess whether ectopic JH could increase it. Therefore, we used the stabilized Geminin context, where endocycling is nearly completely blocked in the ED, to ask whether ectopic JH was sufficient to restore some endocycles in the ED on the day of eclosion. Strikingly, ectopic JH was sufficient to restore significant levels of EdU labeling in EDs expressing stabilized Geminin, with half of samples exhibiting >60 EdU positive cells Figure (8A-C). Since Geminin is a replication licensing inhibitor, this suggests JH acts somehow to either reduce the levels of stabilized Geminin in this context, or to promote DNA replication in parallel to or downstream of replication licensing. We confirmed that treatment with JH did not alter or reduce the levels of *Luna-Gal4* driven *UAS-RFP_nls_* expression suggesting levels of stabilized UAS-driven Geminin should be similar among samples treated with vehicle vs. JH (Supp Fig. 4E). Thus, we favor the hypothesis that JH signaling activates targets in parallel to or downstream of the replication licensing complex to promote endocycling in the adult ED. Indeed, previous work on JH signaling in the female adult fat body of locusts suggests the replication licensing factor *Cdc6* and components of the replication complex *MCMs 4* and *7* are transcriptional targets of JH signaling to promote endocycling (Guo et al., 2014; Wu et al., 2016).

### JH signaling and the endocycle are required for proper protein synthesis in the adult ED

Polyploidy is often correlated with increased biosynthetic capacity for secretory cells (Edgar and Orr-Weaver, 2001). To resolve the biological function of the post eclosion endocycles in the ED driven by JH signaling, we examined whether JH signaling altered the level of nascent protein synthesis in the ED. To test this, we used a well-established puromycin-Click-IT based labeling assay that fluorescently labels nascent proteins (Deliu et al., 2017). When we use *Luna-Gal4* to drive RNAis to *tai*, *Gce* or *met*, we observe significant reductions in protein synthesis in the ED compared to *mCherry* RNAi controls, when normalized to the total area of the ED. (Figure 9A, B). JH signaling has been proposed to promote protein synthesis as well as promote the endocycle, so we next examined whether preventing the post eclosion endocycles could reduce protein synthesis in the adult ED, independent of JH signaling. Indeed, inhibition of the endocycle effectively reduced protein synthesis in the adult ED. We observed significantly reduced nascent protein synthesis in the ED when we knocked down *E2F1* or *E2F2*, or overexpressed *Rbf^280^*, *stabilized Geminin* or *Dacapo*-GFP using *Luna-Gal4* (Figure 9C-F). Protein synthesis is high in the secretory adult ED, but could be further increased by manipulations that increased the endocycling in the ED, such as overexpression of the G1-S *Cyclin/Cdk* complexes *CyclinE/Cdk2* and *CyclinD/Cdk4* using *Luna-Gal4* (Figure 9G, H). These results show that the JH-induced post eclosion endocycling in the ED is required for the proper levels of protein synthesis in the secretory cells of the growing ED.

**Figure 9:**
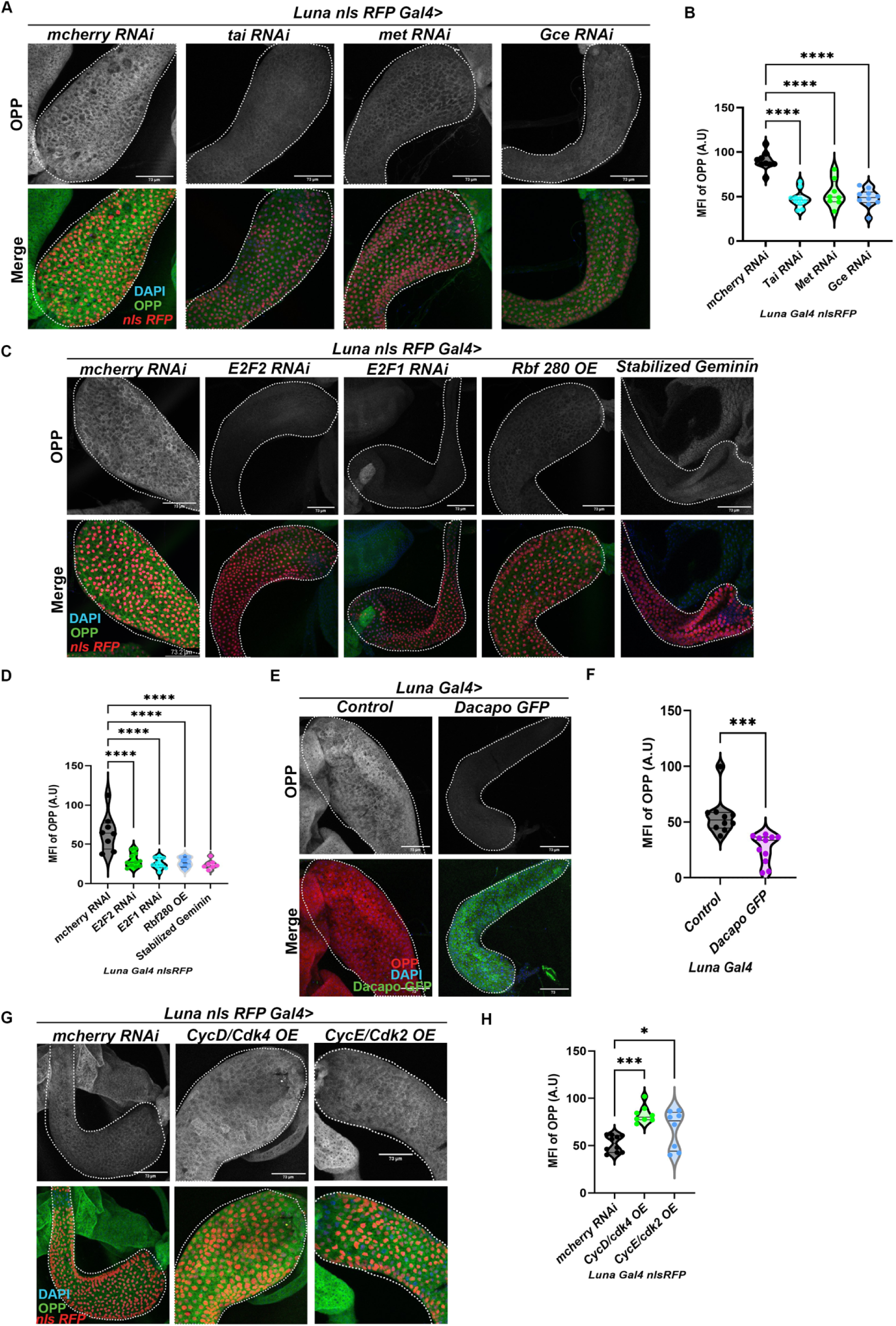
JH signaling and the endocycle are required for proper protein synthesis in the ED. A. Confocal images of ED after OPP nascent protein synthesis assay for 10 minutes on Day 10 with the indicated transgenes driven by *Luna-Gal4, with UAS-nls RFP.* The top panel shows OPP incorporation, the bottom panel is a merge with OPP in Green, *UAS-nlsRFP* in red and DAPI in blue. Scale Bar:73µm B. Quantification of the Mean Fluorescence Intensity of OPP in ED on Day 10 with *mcherry, tai*, *Gce*, *met* RNAi, driven using *LunaGal4nls RFP.* OPP intensity is normalized to the total area of the ED. C. Confocal images of ED after OPP nascent protein synthesis assay for 10 minutes on Day 10 with *mcherry, E2F1* or *E2F2* RNAi, or overexpression of *Rbf280* or stabilized *Geminin* driven using *LunaGal4nls RFP.* The top panel is the images of OPP incorporation, the bottom panel is the merge with OPP in green, *UAS-nlsRFP* in red and DAPI in blue. Scale Bar:73.2µm D. Quantification of the Mean fluorescence intensity of OPP from panel C that is normalized to the area of the ED. E. Confocal images of EDs after OPP nascent protein synthesis assay for 10 minutes on Day 10 with *control* and Overexpression of *dacapo-GFP* using *LunaGal4.* The top panel is the images of OPP incorporation, the bottom panel is the merge with OPP in red, *dacapo-GFP* expression in green, and DAPI in blue. Scale Bar:73µm F. Quantification of the Mean fluorescence intensity of OPP from panel E that is normalized to the area of the ED. G. Confocal images of EDs after OPP nascent protein synthesis assay for 10 minutes on Day 10 with the indicated transgenes driven by *Luna-Gal4 with UAS-nlsRFP.* The top panel is the images of OPP incorporation, the bottom panel is the merge with OPP in green, *nls RFP* expression in red, and DAPI in blue. Scale Bar:73µm H. Quantification of the Mean fluorescence intensity of OPP from panel G that is normalized to the area of the ED. Statistical Analysis: Two-way ANOVA with multiple comparisons between control and Different knockdown, overexpression conditions. P<0.0001 ****

### Protein synthesis in secretory cells of the ED requires mTOR signaling and is coupled to ED growth via the endocycle

The mTOR signaling pathway is a key regulator of protein synthesis (Yang et al., 2022). To test whether protein synthesis in the ED is modulated by mTOR signaling we manipulated components of the pathway, including overexpressing the mTOR activating GTPase *Rheb*, and inhibiting core pathway components such as mTOR and the mTOR Complex 1 (*mTORC1*) target, ribosomal *S6 Kinase* (S6K), by RNAi. Rheb overexpression using *Luna-Gal4* was sufficient to increase protein synthesis in the secretory cells of the ED, without promoting significant additional endocycling or organ growth. (Figure 10 A, B, E, F). By contrast, we observed significantly reduced protein synthesis when we knocked down S6K and mTOR using *Luna-Gal4* (Fig. 10 C,D). We observed a mild reduction in the percentage of 32C cells with the S6K knockdown, although this did not affect organ growth, whereas mTOR knockdown significantly reduced ploidy and ED size (Figure 10E,F). This demonstrates that proper protein synthesis in the secretory cells of the ED requires mTOR signaling and that severe mTOR inhibition can reduce ED endocycling to impact organ growth. We suggest that modulation of mTOR signaling sufficient to alter protein synthesis levels but not the endocycle does not impact organ growth, while severe inhibition of mTOR does reduce endocycling, which reduces organ growth.

**Figure 10:**
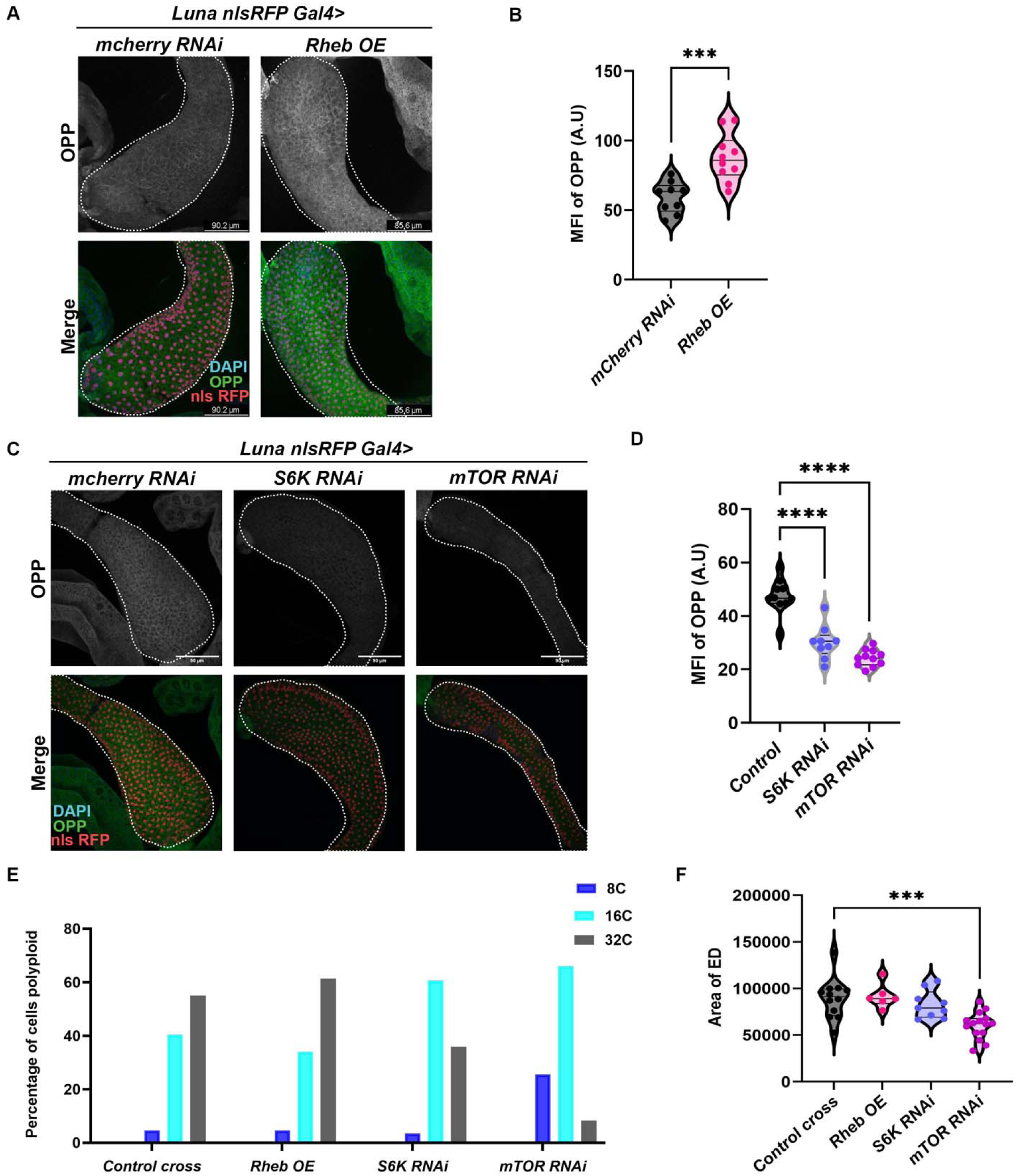
Protein Synthesis in the secretory cells of the ED requires mTOR signaling and is coupled via the endocycle. A. Confocal images of EDs after OPP nascent protein synthesis assay for 10 minutes on Day 10 with *mCherry RNAi* or overexpression of *Rheb* using *LunaGal4* with *UAS-nls RFP.* The top panel is the images of OPP incorporation in the ED, the bottom panel is the merge with OPP in green, *nlsRFP* in red, and DAPI in blue. Scale Bar:90.2µm B. Quantification of the Mean Fluorescence Intensity of the OPP on Day 10 from panel A, OPP intensity is normalized to the area of the ED. C. Confocal images of EDs after OPP nascent protein synthesis assay for 10 minutes on Day 10 with *mCherry, S6K*, or *mTOR* RNAi, driven by *LunaGal4* with *UAS*-*nls RFP.* The top panel is the images of OPP incorporation, the bottom panel is the merge with OPP in green, *nlsRFP* in red, and DAPI in blue. Scale Bar:90µm D. Quantification of the Mean Fluorescence Intensity of the OPP on Day 10 from panel C, OPP intensity is normalized to the area of the ED. E. Quantification of the DNA content/ploidy of the secretory cells of ED on Day 10 post-eclosion for controls expressing *mCherry RNAi*, versus *Rheb* overexpression, knockdown of *S6K* or *mTOR* by RNAi. Reduced ploidy is observed in *S6K* or *mTOR* knockdown conditions. F. Quantification of the area of the ED on Day 10 post-eclosion with overexpression of *Rheb*, or knockdown of *S6K* or *mTOR* with RNAi driven by *Luna Gal4 with UAS-nls RFP*. Statistical Analysis: Figure 10B: Two-tailed unpaired t-test. P<0.0001 ****. Figure 10D, F: Unpaired Two-way ANOVA with multiple comparisons between control and Different knockdown and overexpression conditions. P<0.0001 ****

## Discussion

In this work we demonstrate that the ploidy of the secretory cells of the *Drosophila* male ED impact organ growth and function. Our data suggests this organ is shaped in part through regulated endocycling, which is required for proper protein synthesis levels and organ growth to support male fertility. The growth and endoreplication in the young adult male ED is dependent on Juvenile Hormone signaling post eclosion, and ectopic JH is sufficient to drive additional endocycling in the early adult tissue. The *Drosophila* ED is an important post-mitotic tissue, essential for male fertility, where hormonal signaling, cell growth pathways and cell cycle regulation intersect to impact tissue morphogenesis and function (Figure 11).

**Figure 11.**
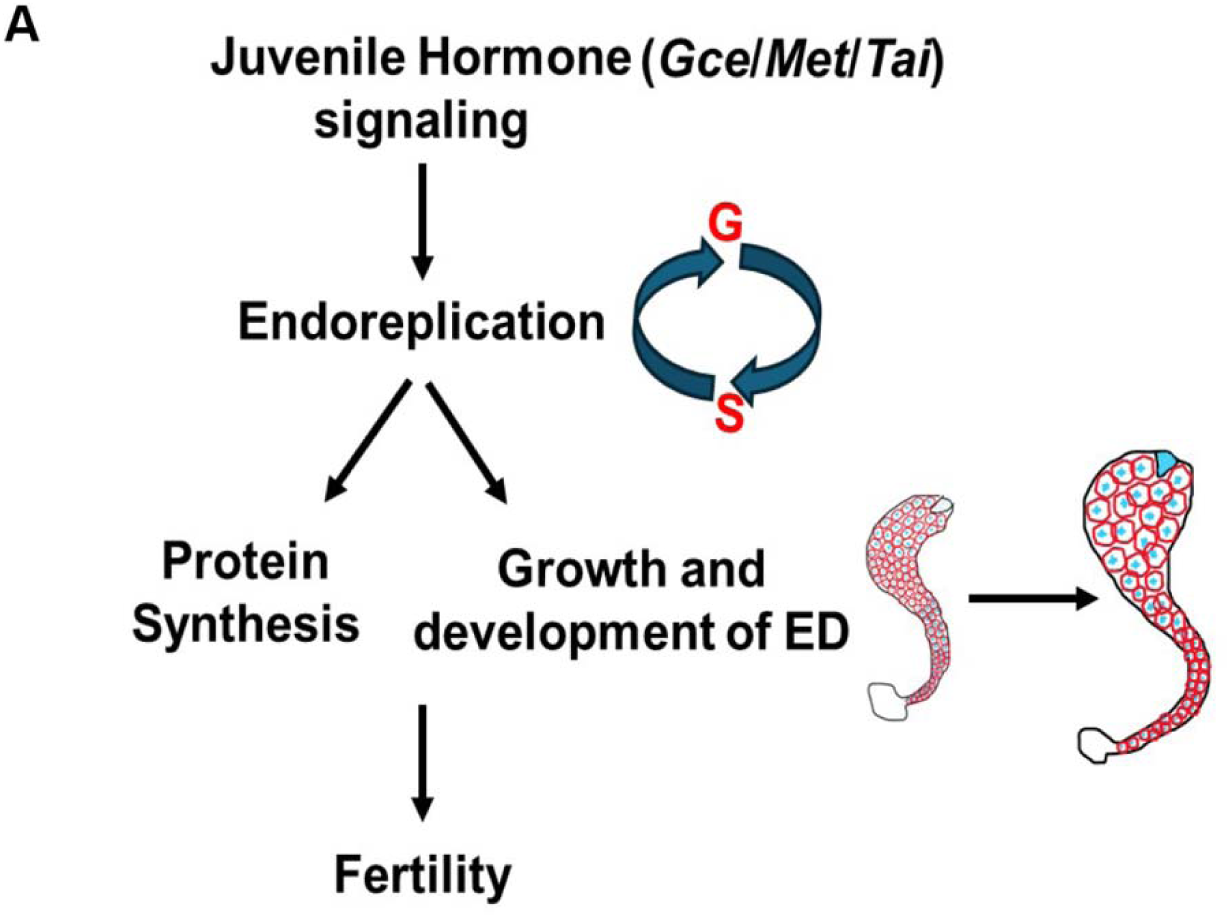
A model for ED organ growth and protein synthesis driven by endoreplication induced by JH signaling.

### Post-eclosion endoreplication in the secretory cells of the ED is required for organ growth, protein synthesis and male fertility

Here we show that the secretory cells of the male ED undergo endoreplication in young adults leading to increases in organ size and ploidy. The cells largely exit from endoreplication by day 3 and reach a maximum ploidy state with the mature organ composed of largely 16C and 32C cells, that are maintained throughout adulthood. When we directly manipulate core cell cycle machinery, ED size, protein synthesis and fertility are affected. While we do not see significant increases in fertility when organ growth is driven through additional endocycling, we do see significant decreases in fertility when endocycles are compromised. Many secretory cells require cell cycle variants that lead to the production of additional copies of genes to support uniquely high biosynthetic requirements for specialized cells (Edgar and Orr-Weaver, 2001). Perhaps the most striking examples are the cell cycle alterations that lead to the amplification of specific genes, such as that observed in the *Drosophila* follicle cells of the ovary preparing to secrete the chorion for the egg (Orr-Weaver, 1991) or the trophoblast giant cells of the mammalian placenta (Hannibal and Baker, 2016). Extensive endoreplication and polyploidization is also seen in the highly secretory cells of the *Drosophila* larval salivary gland, which secrete extensive amounts of glue proteins and components required for pupal adhesion during metamorphosis (Chung et al., 2014). The male *Drosophila* ED, is made up of about 500 cells, yet is required to synthesize a significant amount of seminal fluid components to support male fertility. Our protein synthesis assays in this tissue show that this process is highly active in these cells, modulated by growth and hormonal signaling, and highly responsive to alterations in cellular ploidy.

The normal ploidy distribution in the ED is highly reproducible from animal to animal, suggesting it is tightly regulated. We only observe ploidies higher than 32C with strong direct cell cycle manipulations, such as direct activation of G1-S cell cycle regulators or inhibition of cell cycle checkpoint regulators such as the G1-S inhibitor Rbf or the replication licensing inhibitor Geminin. Even under these strong cell cycle manipulations, the maximum DNA content observed in the ED is 64C across adult timepoints, suggesting the ED cells are only capable of limited extra rounds of endoreplication. Further work will be needed to understand the additional mechanisms promoting cell cycle exit in these cells that limit their ability to endoreplicate further.

### JH promotes the early adult endoreplication in secretory cells of male *Drosophila* ED

JH signaling is involved in multiple aspects of *Drosophila* male fertility (Braun and Wyatt, 1995; Kurogi et al., 2024; Wilson et al., 2003). We recently showed that JH signaling is required for an early post-eclosion endocycle in the adult male *Drosophila* accessory gland (Box et al., 2024) and here we show that JH performs a similar function in the adult ED. JH signaling has been shown to promote polyploidization by activating the transcription of the genes involved in regulating S-phase and DNA synthesis (Guo et al., 2014; Wu et al., 2016; Wu et al., 2020). While we do not yet know the direct targets of JH signaling in the male accessory gland or ED, our finding that JH is sufficient to drive additional endocycling in tissues where the endocycle is blocked by stabilized Geminin, suggests the critical targets are likely to be downstream of the typical G1-S cell cycle regulators and may act at the level of DNA replication licensing and replication. Provocatively, JH receptors Gce and Met have been shown to activate components of the DNA replication complex such as Mcm4 and Mcm7, and the DNA replication licensing factor Cdc6, by binding to an E-box region of their promoters in female locust fat body (Guo et al., 2014; Wu et al., 2016; Wu et al., 2020). Further work will be needed to identify the targets of JH signaling that drive endoreplication in the secretory cells of the male reproductive system.

### Using the *Drosophila* ED as a context to study post-mitotic tissue growth

Studies of the ED have been limited to descriptive approaches such as transcriptomics and proteomics due to the lack of a tissue-specific Gal4 driver to permit inducible genetic manipulations. Here we show that a specific *Luna-Gal4* driver can be effectively used to manipulate pathways regulating hormonal signaling, protein synthesis and the cell cycle in the adult ED. The driver turns on in the late pupal stages a few hours prior to eclosion and maintains expression throughout adulthood, independent of mating status. Using this driver we show that the mTOR signaling pathway intersects with the cell cycle pathway, such that severe inhibition of mTOR signaling leads to a reduction in endocycling and cellular ploidy. This intersection may be through mTOR regulation of *E2F1* protein production (Ovrebo et al., 2022), as *E2F1* is required for proper endocycling in this tissue (Fig 4) and across other endocycling tissues (Zielke et al., 2011). While severe inhibition of mTOR signaling limits endocycling, increased mTOR signaling is sufficient to increase protein synthesis without significantly increasing endocycling in this tissue. Thus, milder manipulations of mTOR signaling allow us to separate out the effects of altering protein synthesis from effects on the endocycle that more strongly impact organ size. JH signaling has been shown to promote both protein synthesis for reproduction and the endocycle (Guo et al., 2014; Wu et al., 2016; Wu et al., 2018; Wu et al., 2020; Yamamoto et al., 1988). Here, because these pathways are linked through mTOR, we cannot rigorously discern whether JH acts through one or both of these pathways in the ED. But our data strongly suggests the impact of JH on the endocycle is likely to be the major effector of its impact on postmitotic ED growth. Additional work will be needed to discern the targets of JH signaling in this tissue, to allow for manipulations of specific outputs of the hormonal growth response.

The similar response of the accessory glands and ED to the post eclosion JH pulse suggests that these two secretory, polyploid tissues are physically linked, as well as linked in function and hormonal response. Since these tissues are involved in producing and secreting seminal fluid components required for male fertility, we suggest that together these tissues can be considered to compose the *Drosophila* prostate-like organ. The mouse prostate is made up of multiple lobes connected to the urethra, while in humans the prostate is made up of multiple zones (Wang et al., 2018). We suggest the *Drosophila* prostate-like organ is made up of two tissues, the accessory glands and the ED, connected via a transition zone made up of the anterior ED papillae. Further work will be needed to examine the cell cycle status of the anterior ED papillae, and we note that accessory gland drivers such as *Paired-Gal4* and the ED driver *Luna-Gal4* described here, do not drive expression in the transition zone.

A major regulator of mammalian prostate growth and function is the steroid hormone Androgen (Banerjee et al., 2018; Massie et al., 2011). Androgen signaling in the mammalian prostate intersects with cell cycle regulation and is a major regulatory axis in mammalian prostate regeneration and cancer (Balk and Knudsen, 2008; Schiewer et al., 2012). Steroid hormone signaling via the ecdysone pathway has been shown to play a role in regulating growth and the endocycle in the male *Drosophila* accessory gland (Leiblich et al., 2019; Sekar et al., 2023; Sharma et al., 2017), but whether ecdysone impacts the ED remains unknown. This question can now be addressed using the ED Gal4 driver we discovered. Our work on JH signaling in the accessory gland has shown that steroid hormone signaling via ecdysone is not the only hormone signaling axis regulating the accessory gland endocycles. Similarly, we expect that JH will not be the only hormonal signaling pathway regulating endocycling and growth in the ED. JH and ecdysone signaling often act in opposition to each other and can converge on shared targets. Future work will be needed to discern whether there is an interplay between JH and ecdysone signaling pathways in these reproductive tissues.

## Materials and Methods

### Drosophila husbandry

All the flies used in this paper were housed at room temperature (23°C) on a Bloomington Cornmeal food without crowding, for all the DOE experiments males were collected immediately post-eclosion (0-4hrs) in a fresh media vial. All the male flies used in all the experiments are virgins except for fertility assays. For day 10 experiments the males were collected on DOE and transferred 5 to 8 flies per vial, aged till day 10 from DOE/Day 1 flipping into fresh media vials at an interval of 2-3 days and housed at room temperature.

### Fly strains used

Canton S

The following *Drosophila* species were kindly provided by the P. Wittkopp Lab (U. Michigan)

*D. simulans*

*D. yakuba*

*D. pseudoobscura*

*D. willistoni*

*D. virilis*

Other lines used for the study are as follows: Luna-Gal4 (BDSC#63746)

Luna-Gal4 recombined with UAS-*nlsRFP* mcherry-RNAi (BDSC #35785)

E2f1-RNAi (BDSC #27564)

E2f2-RNAi (BDSC #36674)

Rbf280 (BDSC #50748)

Dacapo-OE: UAS-Dacapo-GFP (BDSC #83337)

CycE-Cdk2-OE: y,w, hs-flp; +; UAS-CycE,UAS-Cdk2/TM6B (Buttitta et al., 2007) CycD-Cdk4-OE: y,w,hs-flp; UAS CycD, UAS-Cdk4 (Datar et al., 2000)

Geminin-RNAi (BDSC #30929) Rbf-RNAi: (VDRC v10696) Gce-RNAi (BDSC #26323)

Tai-RNAi (BDSC #32885)

Met-RNAi (BDSC #26205)

Stabilized Geminin: UAS-Gem DeltaD/DeltaKEN (BDSC #59069) mTOR-RNAi (BDSC #33951)

Rheb OE: UAS-Rheb on 3 (BDSC #9689) S6K-RNAi (BDSC #41702)

### ED dissection, fixation, and Immunostaining

*Drosophila* male EDs were dissected in 1X Phosphate buffer Saline (PBS), the dissected EDs were immediately transferred to an Eppendorf containing 1ml of 4% Paraformaldehyde (PFA) in 1X PBS to fix the tissues for 30 minutes under continuous rocking. Fixed EDs were rinsed twice using 0.1% Triton-X +1× PBS for 10 minutes each wash. Fixed EDs were permeabilized with 1% Triton-X+1X PBS for 30 minutes on a rocker. Permeabilized tissues were blocked with PAT (1X PBS, 0.1% Triton X-100, 1% BSA) for 10 min. Allow tissues to settle and remove PAT. Resuspended tissues in primary antibody that were freshly diluted in PAT with an appropriate concentration (Mentioned below) and incubated overnight at 4°C on a rocker and washed these tissues with 1XPBS 0.1% Triton X-100, 3 times 10 min. Before the secondary antibody, the tissues were blocked with Normal Goat Serum (1× PBS + 0.1% bovine serum albumin + 0.3% Triton-X (PBT)-X, 2% Normal Goat Serum (NGS)) for 10 min at room temperature. Secondary antibodies was diluted to 1:2000 in NGS and incubated EDs for 4 hours under continuous rocking at room temperature in the secondary antibody. EDs were washed thrice with 1X PBS 0.1% Triton-X after the secondary antibody incubation with 10 minutes each wash. For actin staining we added 2 drops of Actin Red 555 in 1ml of PBS and incubated the EDs for 30 mins and washed twice with 10 mins each wash to remove unbound stain and incubated the EDs in DAPI 1µg/ml for 15 minutes to stain the DNA. EDs were washed with 1× PBS + 0.1% Triton-X after the DAPI staining and mounted in vectashield (Vector Labs). Tissues were mounted by creating a spacer between the slides and coverslip with double-sided tape or one layer of clear nail polish. The list of antibodies used is in the following table with their dilutions.

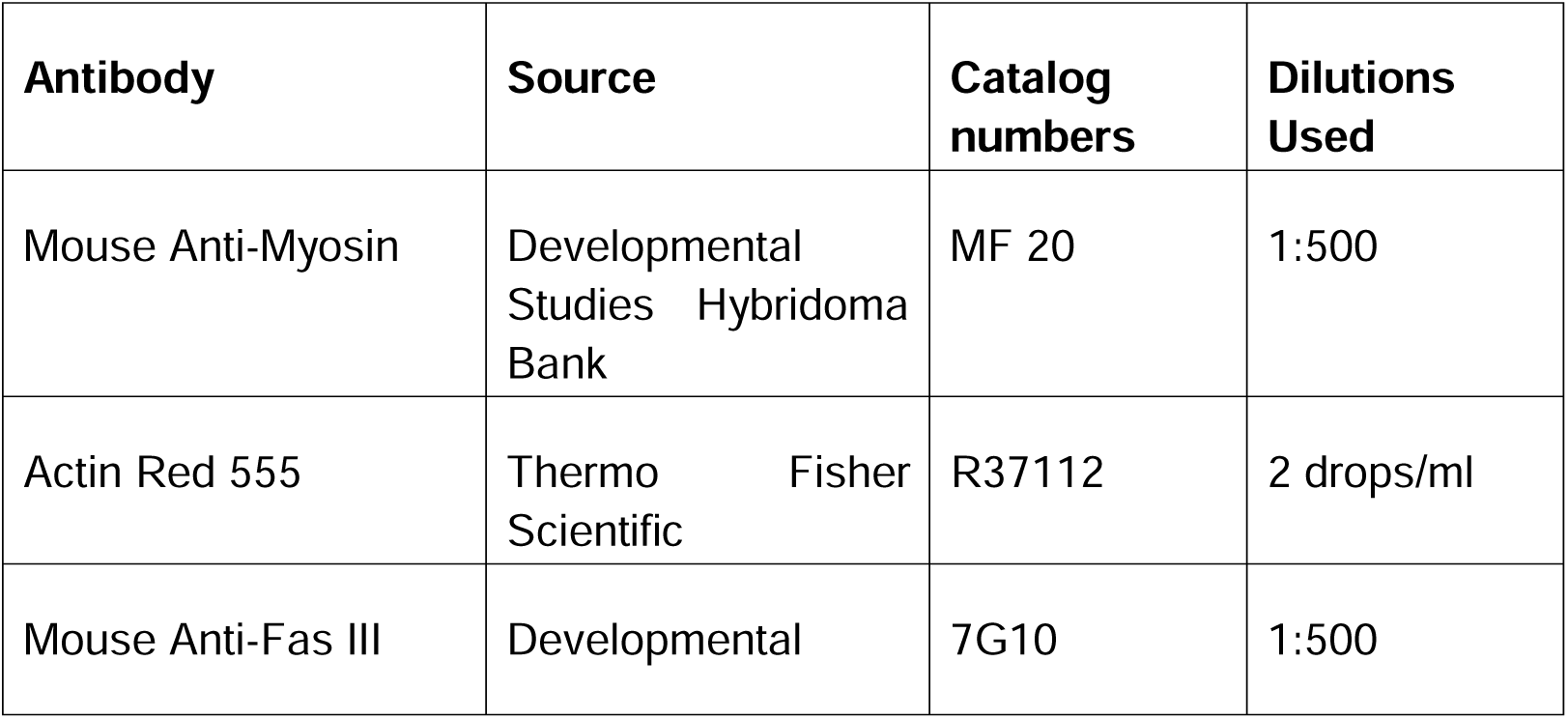
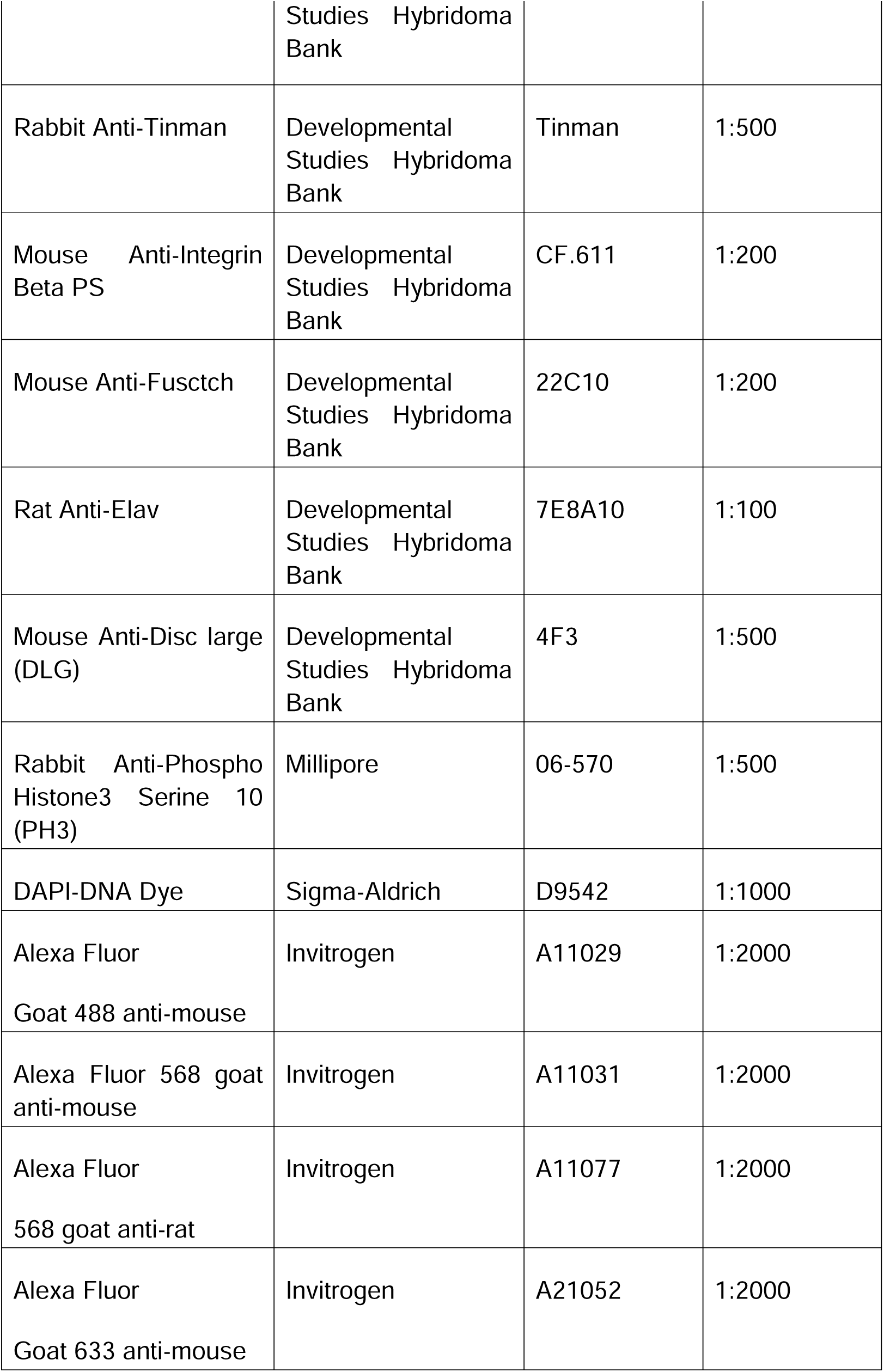

### 5-Ethynyl-2-deoxyuridine (EdU) feeding and labeling

Two different parameters were used to label the S-phase DNA synthesis using EdU. For all the different time point staining of endoreplication on DOE we dissected the EDs in *Drosophila* Ringer’s solution (Sullivan, 2000) and incubated the dissected live EDs in 0.1mM of EdU in Ringers Solution for 1 hour at room temperature (23°C) on a rotator with medium speed. After 1-hour EdU incubation, EDs are fixed, permeabilized and further to visualize the uptake of EdU in the secretory cell’s DNA we performed an iClick-it reaction as mentioned in the reagents protocol. For prolonged labeling of the endoreplication in ED secretory cells, we fed the animals with an EdU solution made with 1mM EdU, 10% Sucrose, and blue food coloring (5µl/ml) to assess the uptake of the EdU by looking at the belly color. In an empty vial, a watchman paper was placed, with the above-made EdU solution (250µl in each vial) in it, flies were allowed to feed on this solution for the mentioned number of days. The Whatman paper was replaced with a new one with a freshly made EdU solution to prevent contamination due to the presence of sucrose. After the feeding time points EDs were dissected and proceeded with fixing, permeabilizing, and iclick-it reaction. We used Click-IT Plus 5-ethynyl-2′-deoxyuridine (EdU) AlexaFluor-555 or 488 (Thermo Fisher C10638) for all the Endoreplication labeling experiments. For analyzing the MFI of the EdU in EDs we imaged the EDs at 10X magnification with keeping the intensity same throughout all the samples. Using imageJ Fiji we quantified the intensity of EdU per unit area of ED.

### Confocal imaging

EDs for all the experiments were imaged using a Leica SP8 Confocal microscope with 40X objective. Some of ED’s 10X and 20X magnification images were imaged using Leica SP5 Confocal microscope and Leica DMI6000B epifluorescence system.

### Area measurements

A complete ED image was captured in one frame using a Leica SP5 Confocal microscope with 10X objective. The images were opened in Fiji Imagej software and obtained the maximum Z projection of the stacks of the image. Manually using the free hand selection tool the area around the ED was drawn from the Anterior end of the ED where the AG, Seminal vesicles are connected to the posterior end of the ED where it connects to the EB. Now the area around the ED was measured by clicking Analyze and then Measure. These area values were plotted using the Graph pad Prism.

### Juvenile Hormone *ex-vivo* assay

Juvenile Hormone III (Sigma Aldrich #J2000) was reconstituted in Acetone to make the stock concentration of 10mg/ml. We used a concentration of 0.1µg/ml in Ringer’s solution for all the JH *ex-vivo* experiments. The EDs were dissected in Ringer’s solution immediately after eclosion and incubated in JH III along with 0.1mM of EdU to assess the endoreplication levels in the presence and absence of JH III for 1 hour under continuous rotation at room temperature. The control cross was also incubated with the same concentration of JH and EdU. As a vehicle control, we added just Acetone in Ringer’s solution and EdU in a separate watch glass for both the control cross and the test. After the one-hour incubation of JH and EdU the tissues were fixed, permeabilized, and stained.

### Ploidy/DAPI intensity measurements

All the ED DAPI images were obtained without saturating the DAPI intensity using a Leica SP8 confocal microscope. Images were exported in Tiff format and opened in Fiji Imagej. The maximum projection of the stacks of DAPI-stained images was obtained. We have observed that the basally located nuclei in the external muscle layer of the ED are diploid by flow cytometry. We used these diploid muscle nuclei of ED to normalize the measurements and to calculate the haploid nuclei intensity. We measured the background intensity by drawing the region of interest (ROI) in the background region of the images. After the background subtraction ROI was manually drawn around the DAPI-stained nuclei of Diploid and polyploid secretory cells. Measured the Raw integrated Intensity of both diploid and secretory cells polyploid nuclei. We measured around 30 secretory cell polyploid nuclei and 10 diploid nuclei from muscle for each ED and We calculated the corrected fluorescence intensity from each nucleus raw integrated intensity, by subtracting the average background intensity. The intensity of the haploid nuclei DNA was calculated by taking the average of the diploid nuclei intensity and dividing it by half, we used this as a reference of haploid nuclei with DNA content of 1C. Based on these haploid nuclei we estimated the DNA content of polyploid secretory cell nuclei. For Figures 4F we directly scored the DNA content without binning. For all other DNA content/ploidy measurements we plotted the after the following binning-2N (1.9-2.9), 4N (3.0-6.9), 8N (7-12.9), 16N (13.0-24.9), 32N (>24.9).

### Fertility Assay

Virgin males were used to set up the fertility assay in all the genotypes mentioned in the figures. We used 2-3 day old virgin Canton S females to cross the males of each genotype. We used a ratio of 1:8 (Male: Female) to set up the fertility assay in each vial and allowed them to mate for 3 days in 25°C incubator. The total number of the pupae and the offspring eclosed were counted and plotted as the result of fertility.

### Flow cytometry

Nuclei from the ED secretory cells were isolated at the mentioned time points. We used S2 media (Thermofisher #21720024) with 10%Fetal Bovine Serum (Cytiva # SH30396.02HI) and 1% Pen-strep antibiotic solution. The EDs were dissected in Ice-cold S2 media+10%FBS+1% Pen-strep. We used 25-30 EDs for the nuclei extraction per sample. We briefly spun the tissues to allow it to settle, S2 media was removed, added 1ml of Lysis Buffer (10mM Tris HCl-pH 7.4, 10mM NaCl, 3mM MgCl_2_, 0.1% NP40) and transferred the tissue with the buffer to Dounce homogenizers that are placed on ice. To release the nuclei we homogenized the tissues with 20 strokes of loose Dounce pestle and allowed the tissue to rest on Ice for 5 minutes. Following this we homogenized with tight Dounce pestle for 40 strokes. Triturated the tissue 15 times with a silanized fire-polished glass Pasteur pipette (BrainBits). This homogenate was filtered through 100 microns filter mesh, followed by 40 micron filter mesh to remove debris and unlysed tissue. To this homogenate 100µl of S2 media was added to stop the lysis reaction. This filtrate was centrifuged at 800 rcf for eight minutes at 4°C. Removed the supernatant and added 200µl of S2 media to the nuclei pellet with and incubated on ice for 5 minutes. To this additional 800µl of S2 media was added and centrifuged for eight minutes at 800 rcf, 4°C, supernatant was removed and the pellet was resuspended in 200µl S2 media with Vybrant DyeCyle Violet (Invitrogen #V35003) at a ratio of 1:100 and allowed to incubate on ice for 5 minutes, after the incubation to this 800µl of S2 media was added and centrifuged at 800rcf for eight minutes at 4°C. The supernatant was removed and resuspended the pellet in 1ml of S2 media, gently vortexed, and run on the Attune Flow Cytometer. We used Ovaries as a positive control to overlay the ED peaks. We followed the same steps to isolate the nuclei from the ovaries and stained them with the same protocol. For flow cytometry on Drosophila species, we overlaid the ED ploidy plot with ovaries from the matching species.

### Protein synthesis Puromycin assay (OPP) Assay

We performed the OPP assay in S2 Media (invitrogen) that has been reconstituted with 10%FBS and 1% Pen-strep. Adult fly ED was dissected in S2 media and incubated with 20 µM of OPP (O-Propargyl Puromycin, Thermofisher) in S2 media for 10 minutes. The tissues were immediately fixed in 4% paraformaldehyde made with 1XPBS for 30 minutes. Fixed EDs were rinsed twice using 0.1% Triton-X +1× PBS for 10 minutes each wash. Fixed EDs were permeabilized with 1% Triton-X+1X PBS for 30 minutes on a rocker. Performed the click-it reaction to label the OPP using iClick-it OPP Kit (Thermofisher #C10456) by following the instructions provided by the kit. The tissues were washed twice with 0.1% Triton-X-1XPBS for 10 minutes each wash, mounted the tissues on the glass slides using vectashield as a mounting medium.

### Software and data analysis

We used Fiji ImageJ to analyze all the data. We used GraphPad Prism software to perform statistical significance in each graph plotted in the figures. The tests used to run the significance are mentioned in the figure legends.

## Acknowledgments

We thank Dr. P. Wittkopp and the Wittkopp Lab for providing *Drosophila* of various species. We thank S.J. Church for contributions to developing the flow cytometry protocol for the ED. We thank Buttitta Lab members for valuable discussions and suggestions on this project. Funding for this work was provided by the National Cancer Institute (P01CA093900) and the National Institute of General Medical Sciences of the National Institutes of Health (R01GM127367 and R35GM149273). A. Box was supported in part by the University of Michigan Organogenesis Predoctoral Training Grant (NIH T32 HD007505) and the Eunice Kennedy Shriver National Institute of Child Health and Human Development of the National Institutes of Health (NIH F31HD103430). The authors have no conflicts of interest to disclose.

