## Supplement for "Post-eclosion growth in the *Drosophila* Ejaculatory Duct is driven by Juvenile Hormone signaling and is essential for male fertility"

**Supplement to Ramesh et al 2024**

**Includes 4 figures with Figure legends**

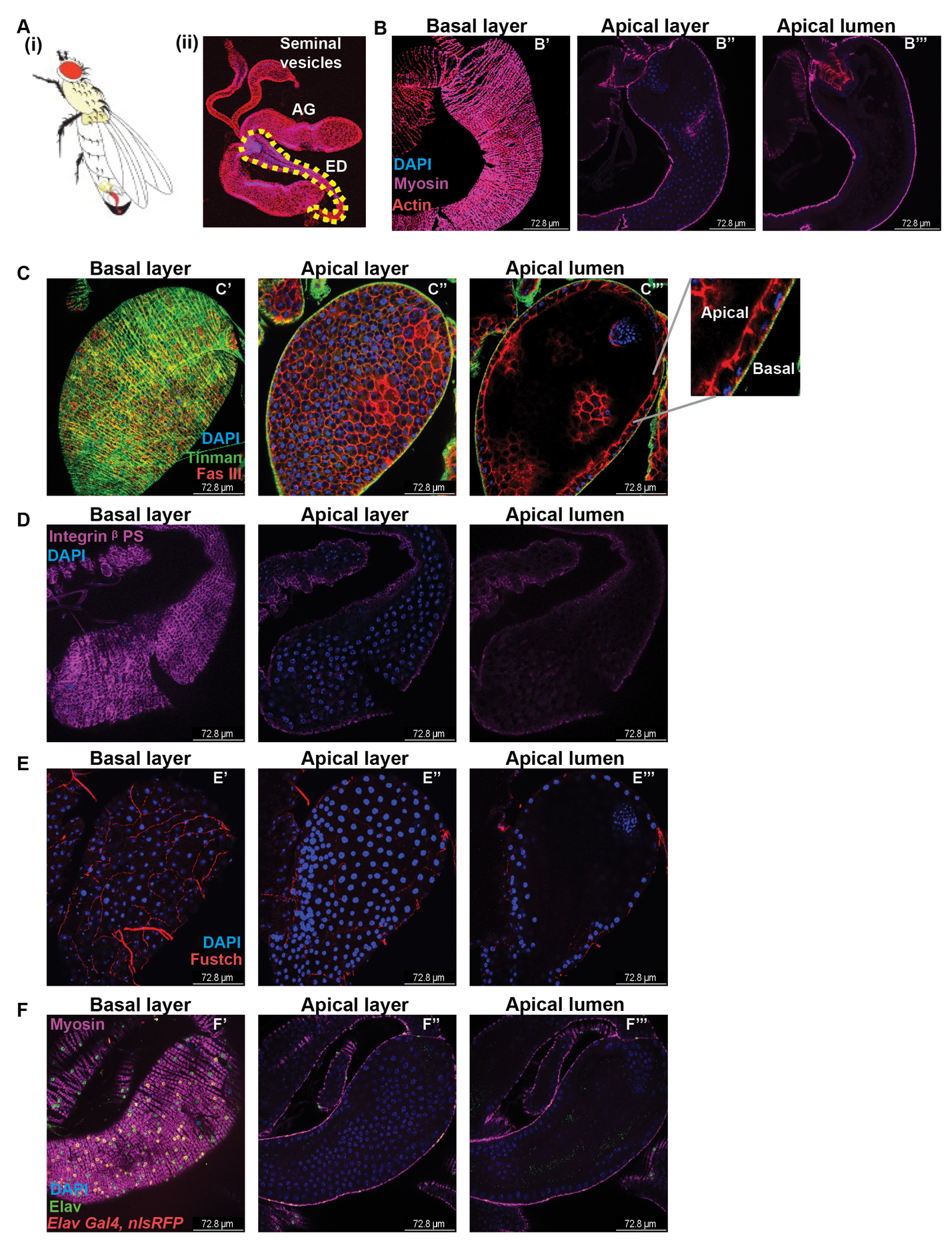

**Supplementary Figure 1: Identification of different cell types in adult *Drosophila* Ejaculatory duct**

A. (i) Cartoon of the *Drosophila* male reproductive system (ii) Micrograph of male reproductive system stained with cell junction marker DLG and nuclear staining DAPI with yellow dotted line highlighting the ED.
B. Confocal micrographs of Myosin, actin, and DAPI staining in different apico-basal planes of the ED, B’ is the outer basal layer, B’’ is the inner Apical layer, and B’’’ is the Inner apical layer of cells that is exposed to the hollow lumen.
C. Confocal micrographs of Tinman staining in the basal layer of ED (C’) and the Apical secretory cells are labeled with cell junction protein Fas III (C’’) with nuclei labeled with DAPI. Zoomed in image of C’’’ showing the apical and basal side of ED.
D. Confocal micrographs of ED stained with Integrin βps antibody which labels an extracellular matrix receptor, and Dapi staining in the apical secretory cells.
E. Confocal micrographs of ED Basal (E’), Apical (E”), and Apical lumen (E’’’) showing the axonal projections that are stained with the Fustch antibody, and nuclei labeled with DAPI.
F. Confocal micrographs of ED showing the Elav staining (green), and the same cells labeled with Elav Gal4 expressing nlsRFP. Muscles are stained with anti-Myosin (magenta). The staining shows that the ED muscles are innervated with neurons and are present on the exterior basal layer. The nuclei are labeled with DAPI. Scale bar: 72.8 microns

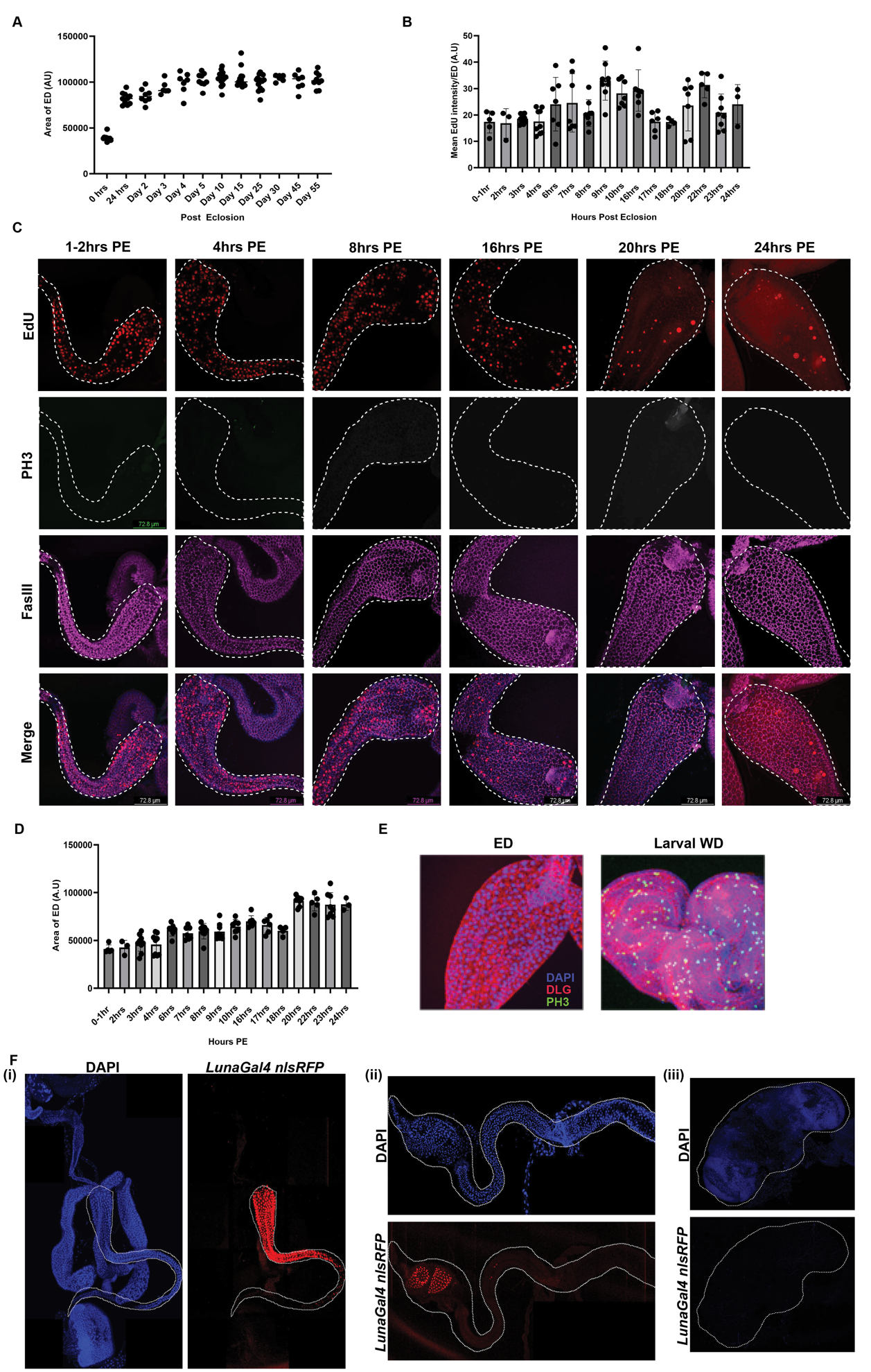

**Supplementary Figure 2**

A. Area of adult EDs at indicated time points post-eclosion. The ED grows from Day of eclosion (DOE) to Day 5 with growth ceasing post day 5. Each point is the measurement of the area of one ED.
B. Quantification of the intensity of EdU labeling in EDs exposed to EdU for one hour intervals post eclosion up to 24hrs. The intensity is normalized to the area of the ED at each time point.
C. Confocal images of EDs labeled with EdU at 1-2hrs, 4hrs, 8hrs, 16hrs, 20hrs, 24hrs post-eclosion performed with *ex vivo* incubation. Top panel in red is the EdU incorporation in ED, next is Phospho histone H3 (PH3) staining in green showing no mitotic cells, next panel is secretory cell junctions labeled with FasIII in Magenta. The bottom panel is the merge of EdU (red), PH3 (green) FasIII (magenta), and DAPI (blue). Scale Bar:50µm
D. Measurement of the area of ED on at an interval of every hour post-eclosion to 24 hrs. ED area increases until 24 hrs post eclosion.
E. Confocal images of EDs and larval imaginal discs stained with PH3 in green, with cell junctions labeled by Discs Large (DLG) (red), and nuclei stained by DAPI (blue).
F. Testing *Luna Gal4* expression in different tissues using *UAS-nls RFP* with nuclei counter-stained with DAPI (i) Confocal image of male reproductive system with *Luna Gal4* expression in only ED. (ii) Confocal image of the gastrointestinal tract with *Luna-Gal4* expression only in the cardia. (iii) Confocal images of the brain showing the absence of *Luna Gal4* expression. Scale bar:333.3 µm.

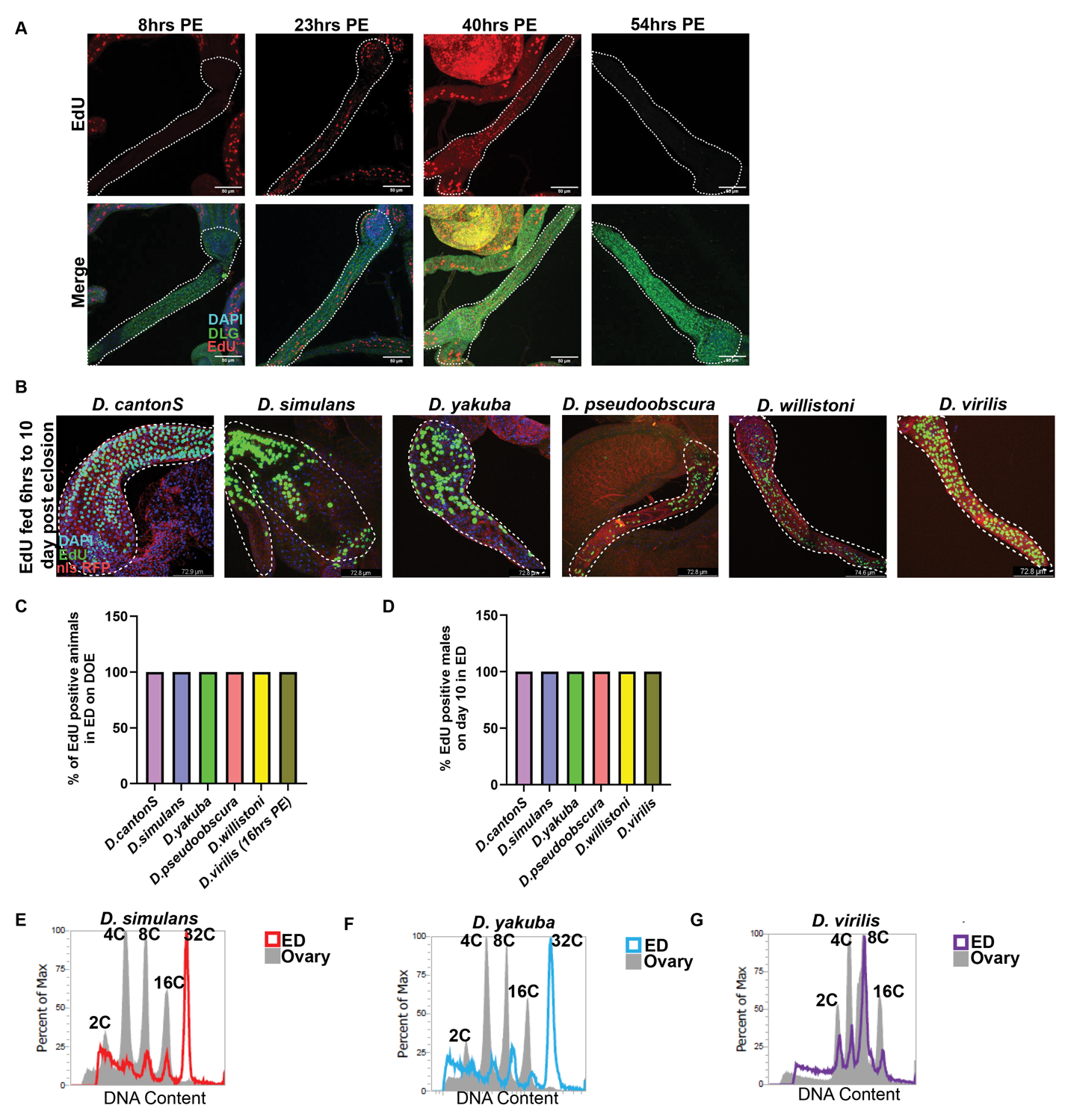

**Supplementary Figure 3**

A. Confocal images of EdU labeling in *D.virilis* to identify the time point when the ED starts endocycling post-eclosion (PE) by soaking the tissue *ex vivo* in EdU for 1hr. Endocycling is absent at 8hrs PE but starts by 16hrs PE, continues for 40-50hrs PE and cell cycle exit occurs by 60 hrs. The top panel is EdU (red) labeling and the bottom panel is the merged image of DAPI (blue), and cell junction marker Discs Large (DLG, green).
B. Confocal images of different species of *Drosophila* fed with EdU from 6hrs to Day 10 PE and dissected on Day 10 and imaged to observe the EdU labeling in secretory cells of ED. DLG (red), EdU (green), DAPI (blue).
C. Quantification of the EdU labeling on day of eclosion (DOE) in different *Drosophila* species. Data shown as the percentage of animals that are positively labeled with EdU during a 1 hour *ex vivo* labeling on DOE.
D. Quantification of EdU labeling on Day 10 in different *Drosophila* species. Data shown as the percentage of animals that are positively labeled with EdU over a 10 day feeding.
E. -G: Flow cytometry histograms of nuclear DNA content of ED secretory cells on Day 10 in *D. simulans, yakuba and virilis* species respectively. Gray shaded peaks in all the three plots are species-matched ovary nuclei to assign correct ploidy peaks for each species genome size.

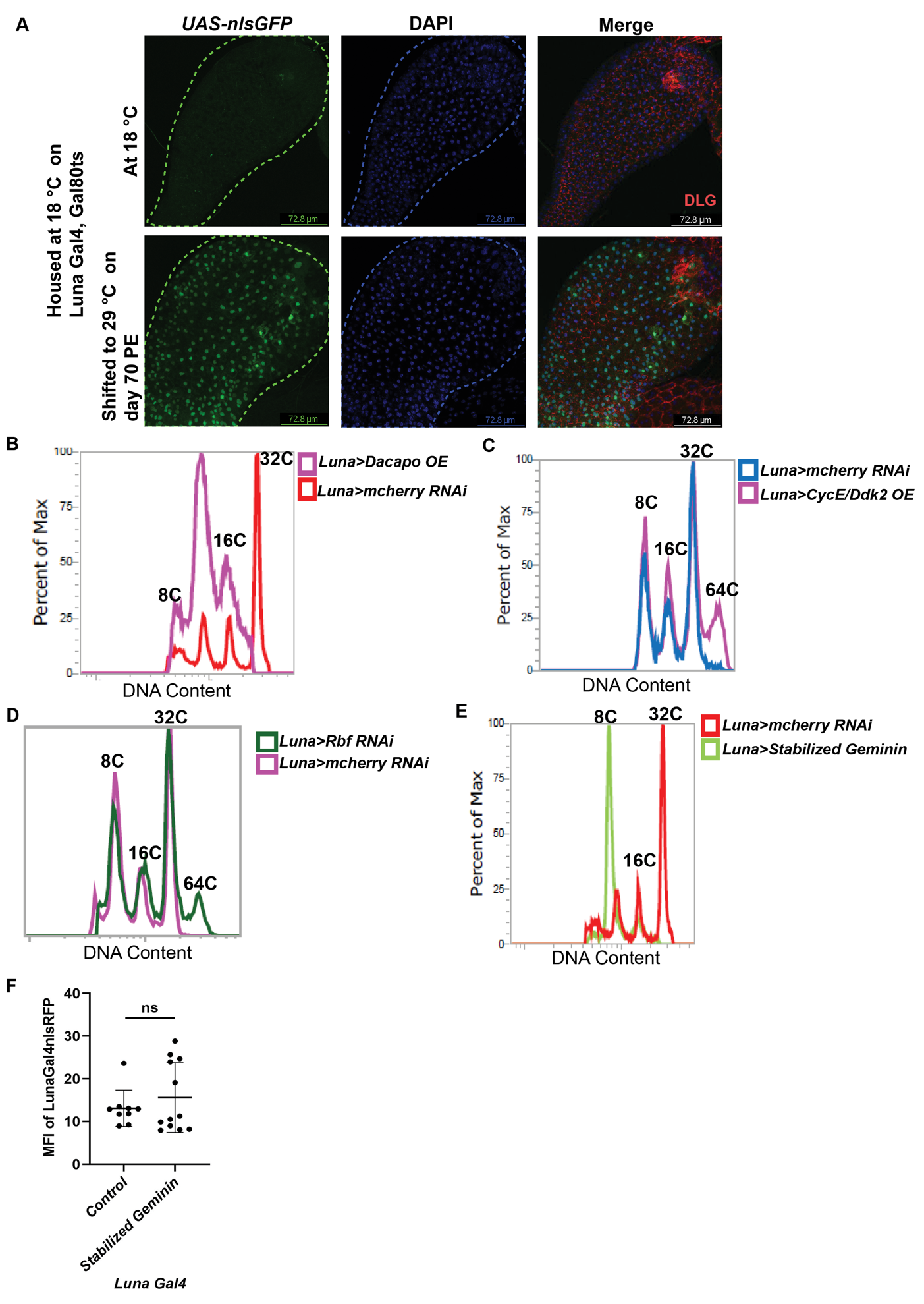

**Supplementary Figure 4**

A. Confocal images of ED showing the expression of *Luna Gal4* on Day 70 post-eclosion (PE) using *UAS-nls RFP.* Scal bar: 72.8µm.
B. Flow cytometry histogram of nuclear DNA content of ED secretory cells on Day 10 with *dacapo-GFP* Overexpression and control *mcherry RNAi.*
C. Flow cytometry histogram of nuclear DNA content of ED secretory cells on Day 10 with *CycE/Cdk2* overexpression vs. control *mcherry RNAi.*
D. Flow cytometry histogram of nuclear DNA content of ED secretory cells on Day 10 with *Rbf RNAi* vs. control *mcherry RNAi.*
E. Flow cytometry histogram of nuclear DNA content of ED secretory cells on Day 10 with stabilized *Geminin* overexpression vs. control *mcherry RNAi.*
F. Quantification of Mean Fluorescence intensity of *UAS-nls RFP* expression in ED driven by *Luna Gal4* in a control cross without an additional transgene vs. co-expression of stabilized *Geminin*
